# Germline mitophagy selectively eliminates deleterious mitochondrial DNA across generations

**DOI:** 10.64898/2026.06.24.734151

**Authors:** Arnaud Ahier, Tessa Onraet, Daniel Campbell, Brendan Townsend, Ayenachew Bezawork-Geleta, Sasheen Dowlath, Anne Hahn, Rachel Shin Yie Lee, Chuan-Yang Dai, Arnaud Gaudin, Julia Pagan, Steven Zuryn

## Abstract

Faithful transmission of genetic information through the immortal germline is essential for organismal health and species survival, yet how mutant mitochondrial genomes (mtDNAs) are selectively eliminated across generations remains unclear. Here, we show that germline mitophagy functions as a mutation-responsive surveillance system that selectively eliminates mutant mtDNAs and shapes inheritance across generations. In *C. elegans*, mitochondria enriched for mutant mtDNAs are selectively removed in the maternal germline prior to oocyte fertilization via PINK1/Parkin-dependent and BNIP3-mediated mitophagy pathways activated by mtDNA defects. Germline mitophagy declines with age, resulting in offspring that inherit increased burdens of mutant mtDNAs. Conversely, enhancing mitophagy within germ cells restricts the transmission of deleterious genomes in a compounding manner, ultimately driving their complete elimination from the matrilineal lineage. Together, our findings demonstrate that germline mitophagy is a critical determinant of intergenerational mitochondrial genome inheritance, establishing its role in restricting the transmission of defective genetic information.

## Introduction

For all obligate aerobes, mitochondria are essential organelles that sustain cellular energy production, metabolism, and signaling, and are indispensable for organismal health, development, and reproduction. Central to mitochondrial function is the mitochondrial genome (mtDNA), which resides within the mitochondrial matrix, encodes core components of the oxidative phosphorylation machinery, and must be faithfully transmitted to progeny to sustain life. Yet the mitochondrial genome presents a fundamental biological challenge: it is maternally inherited, largely non-recombinant, and accumulates mutations at rates far exceeding those of the nuclear genome^1^. These features limit classical genetic mechanisms for error correction and raise a long-standing question – how mitochondrial genome integrity is preserved across generations.

Despite its vulnerability to mutation, mtDNA is transmitted with remarkable fidelity. Evidence across multiple species, including humans, indicates strong purifying selection that limits the inheritance of most deleterious mtDNA mutations, such that relatively few individuals are born with pathogenic variants at levels sufficient to cause disease^2–6^. The genetic bottleneck model proposes that, during oogenesis, a reduced and largely stochastic subset of mtDNA molecules is transmitted from primordial germ cells (PGCs) to mature oocytes, amplifying differences in mtDNA composition between cells and creating conditions under which selection can occur^7–9^. However, the cellular and molecular mechanisms that execute this selection remain poorly defined.

Several mechanisms have been proposed to explain how mutant mtDNAs are selectively removed at the organelle level, including biased mitochondrial transport, replication competition between organelles, and targeted degradation^1,10–13^. Components of mitochondrial stress signaling, such as the mitochondrial kinase PTEN-induced putative kinase 1 (PINK1), which regulates mitochondrial dynamics^14^, local translation^10^, calcium handling^15^, mitochondrial-derived vesicles^16,17^, and mitophagy^18^, have been implicated in restricting mutant mtDNA propagation; however, its function has remained mechanistically ambiguous and canonical downstream machinery of mitophagy has not been consistently linked to mtDNA selection across generations^10,11^. Indeed, in several systems, key mitophagy factors and core autophagy proteins appear dispensable for mtDNA purifying selection, and thus direct evidence that canonical mitophagy actively surveils and removes mutant mtDNAs from the germline and prevents their propagation across generations remains unresolved^10,12,13,18–23^.

Here, we show that canonical mitophagy operates as a selective, mutation-responsive surveillance mechanism within the germline that governs mitochondrial genome quality across generations. Using *C. elegans*, we demonstrate that mechanistically independent PINK-1 (PINK1)-PDR-1 (Parkin) and DCT-1 (BNIP3) mitophagy pathways operate in parallel within germline stem cells (GSCs) to promote the selective autophagosomal engulfment and elimination of mitochondria enriched for mutant mtDNAs. These pathways perceive, respond, and limit the incorporation of mutant genomes into unfertilized oocytes, progressively reduce mtDNA heteroplasmy across generations, and safeguard intergenerational mitochondrial genome integrity. We show that age-dependent or genetic attenuation of mitophagy increases the burden of mutant mtDNAs transmitted to offspring, whereas enhancing mitophagy in the germline rapidly drives their complete elimination from matrilineal lineages, with corresponding improvements in fitness. Together, these findings establish mitophagy as a critical germline quality-control checkpoint that ensures faithful mtDNA inheritance across many generations.

## Results

### Mutant mtDNAs Trigger Autophagosomal Engulfment of Mitochondria in Germ Cells

We first assessed mitophagy activity in the maternal germline of *C. elegans* by examining the recruitment of autophagosomal membranes to mitochondria in the germ cells of live animals. To visualize this process, we generated transgenic strains co-expressing fluorophore-tagged LGG-1, the *C. elegans* ortholog of LC3 that labels autophagosomal membranes and interacts with mitophagy receptors on the mitochondrial outer membrane^24^, together with a germline-specific mitochondrial marker. Colocalization of LGG-1-positive membranes with mitochondria indicated the formation of mitophagosomes (MPs) within the germline (Fig. 1a,b).

**Fig. 1:**
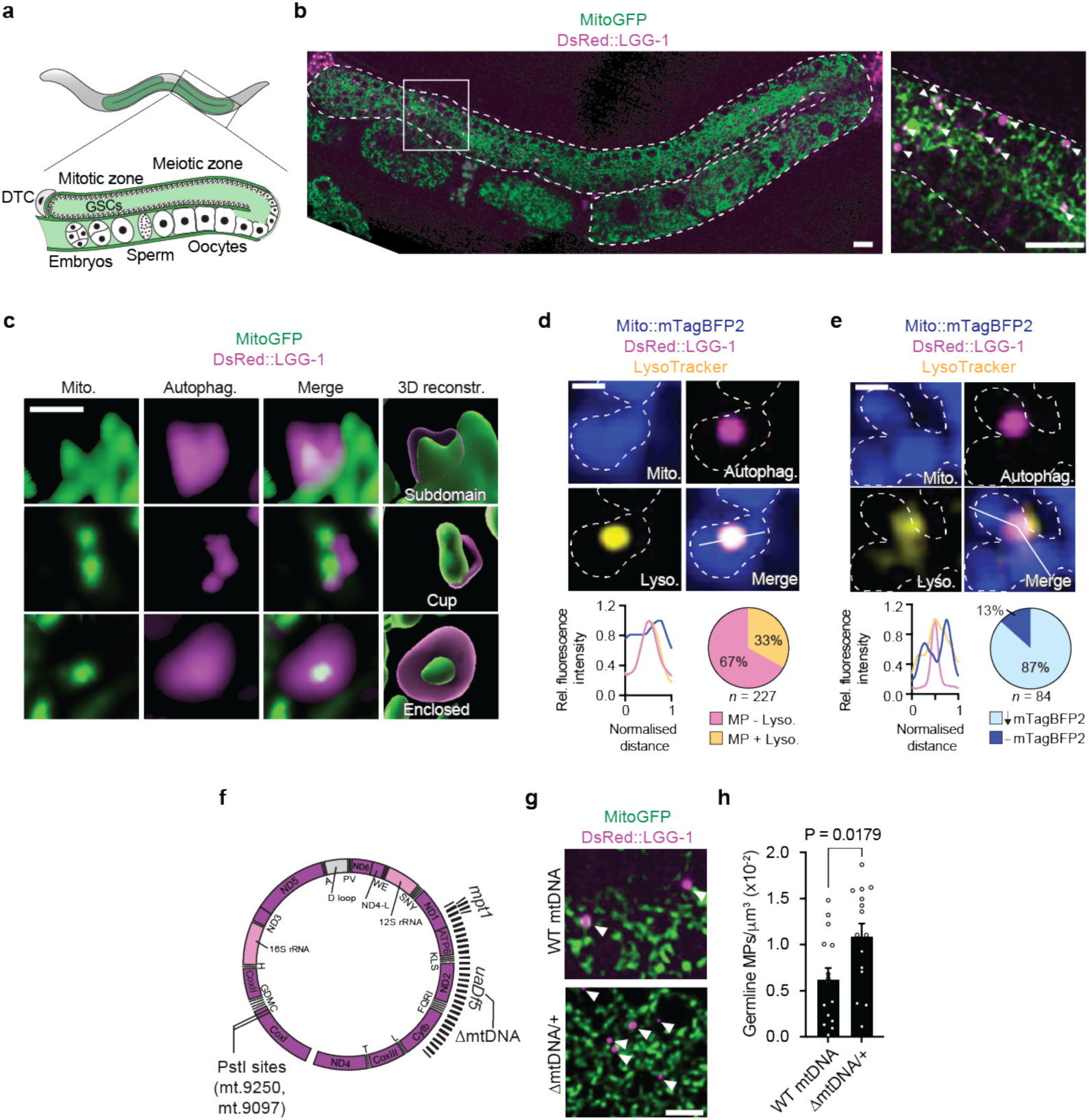
Defective mtDNAs Trigger Autophagosomal Engulfment of Mitochondria in the Germline. **a**, Graphical representation of the *C. elegans* adult hermaphrodite gonad and germline. One of the two gonad arms is shown in detail. Distal tip cells (DTCs) establish a functional stem cell niche that maintains germline stem cells (GSCs). GSCs divide in the mitotic region before entering the meiotic region and progressively developing into oocytes. Mature oocytes are subsequently self-fertilized by sperm to give rise to embryos. **b,** Representative images of mitophagosomes (MPs) in the germline. Left panel, a whole gonadal arm of an adult *C. elegans* hermaphrodite, outlined with a white dashed line, is shown. MPs are defined by the spatial overlap between the autophagosome marker DsRed::LGG-1 (magenta) and a mitochondrial GFP marker (MitoGFP, green). Right panel, higher magnification of the region indicated by the white square. MPs are highlighted with white arrowheads. Scale bar, 10 µm. **c,** Representative images and three-dimensional reconstructions of MPs in the *C. elegans* germline. Upper panel, a subdomain of larger mitochondria in contact with and partially surrounded by a phagophore. Middle panel, smaller mitochondria associated with a cup-shaped phagophore. Lower panel, a small mitochondrial particle fully enclosed by a phagophore. Scale bar, 1 µm. **d,** Representative image of an MP colocalizing with LysoTracker to form an autolysosome-like structure in the germline. Scale bar, 1 µm. Bottom left, fluorescence intensity profile of a scan line traced across the autolysosome-like structure (white line in image). Bottom right, quantification of germline MPs colocalizing with LysoTracker (+ Lyso). **e,** Representative image of an MP colocalizing with a lysosome to form an autolysosome-like structure in the germline, associated with reduced mTagBFP2 mitochondrial signal. Scale bar, 1 µm. Bottom left, fluorescence intensity profile of a scan line traced across the autolysosome-like structure (white line in image). Bottom right, quantification of autolysosome-like structures associated with reduced mTagBFP2 signal (τmTagBFP2). **f,** Representation of *C. elegans* mtDNA showing the location of the 3.1 kb *uaDf5* deletion (ΔmtDNA), the 179 bp *mpt1* deletion, and the PstI restriction sites (mt.9250 and mt.9097). **g,** Representative images of MPs in the germline in the presence or absence of ΔmtDNA. MPs are indicated with white arrowheads. Scale bar, 5 µm. **h,** Quantification of germline MP abundance normalized to mitochondrial volume in the germline in the presence or absence of ΔmtDNA. Columns are mean ± SEM; n = 15 germlines per condition (3 independent experiments); unpaired two-tailed Student’s t-test.

High-resolution imaging captured multiple stages of this process, including phagophores partially surrounding mitochondrial subdomains and mitochondria fully enclosed within autophagosomes (Fig. 1c), consistent with a continuum of mitophagy intermediates. A subset of MPs colocalized with LysoTracker-positive compartments (Fig. 1d), indicating delivery to acidified vesicles. Of these, most exhibited reduced mitochondrial fluorescence (Fig. 1e), consistent with lysosomal degradation or quenching of mitochondrial contents.

We next asked whether mtDNA defects trigger this response, as might be expected if germline mitophagy functions to eliminate mutant genomes. Animals carrying a 3.1-kb mtDNA deletion *(uaDf5*; *Δ*mtDNA) (Fig. 1f), a second deleterious mtDNA allele (*mpt1*), or elevated rates of *de novo* mtDNA mutation caused by inactivation of the proofreading domain of the mitochondrial DNA polymerase γ, POLG-1^25^, exhibited a marked increase in MP formation (Fig. 1g,h and Extended Data Fig. 1a-e), demonstrating that mitophagy is activated by the presence of distinct mtDNA mutations in the germline.

### Germline Mitophagosomes (MPs) Selectively Target Mutant mtDNAs

Because the capacity of mitophagy to eliminate mutant mitochondrial genomes depends on the incorporation of mtDNAs into mitophagic cargo, we examined the content and properties of germline MPs. To visualize nucleoid inclusion, we fluorescently labelled endogenous HMG-5, the *C. elegans* ortholog of the mtDNA packaging nucleoid factor TFAM, using CRISPR-Cas9 genome editing, which revealed individual nucleoids within the majority of germline MPs (Fig. 2a,b and Extended Data Fig. 2a). To further substantiate this, we purified germline autophagosomes from populations of animals by immunoprecipitating HA-tagged LGG-1 that we expressed selectively within their germlines (Fig. 2c). Quantitative PCR analysis of the isolated fractions confirmed enrichment of mtDNA (Fig. 2d).

**Fig. 2:**
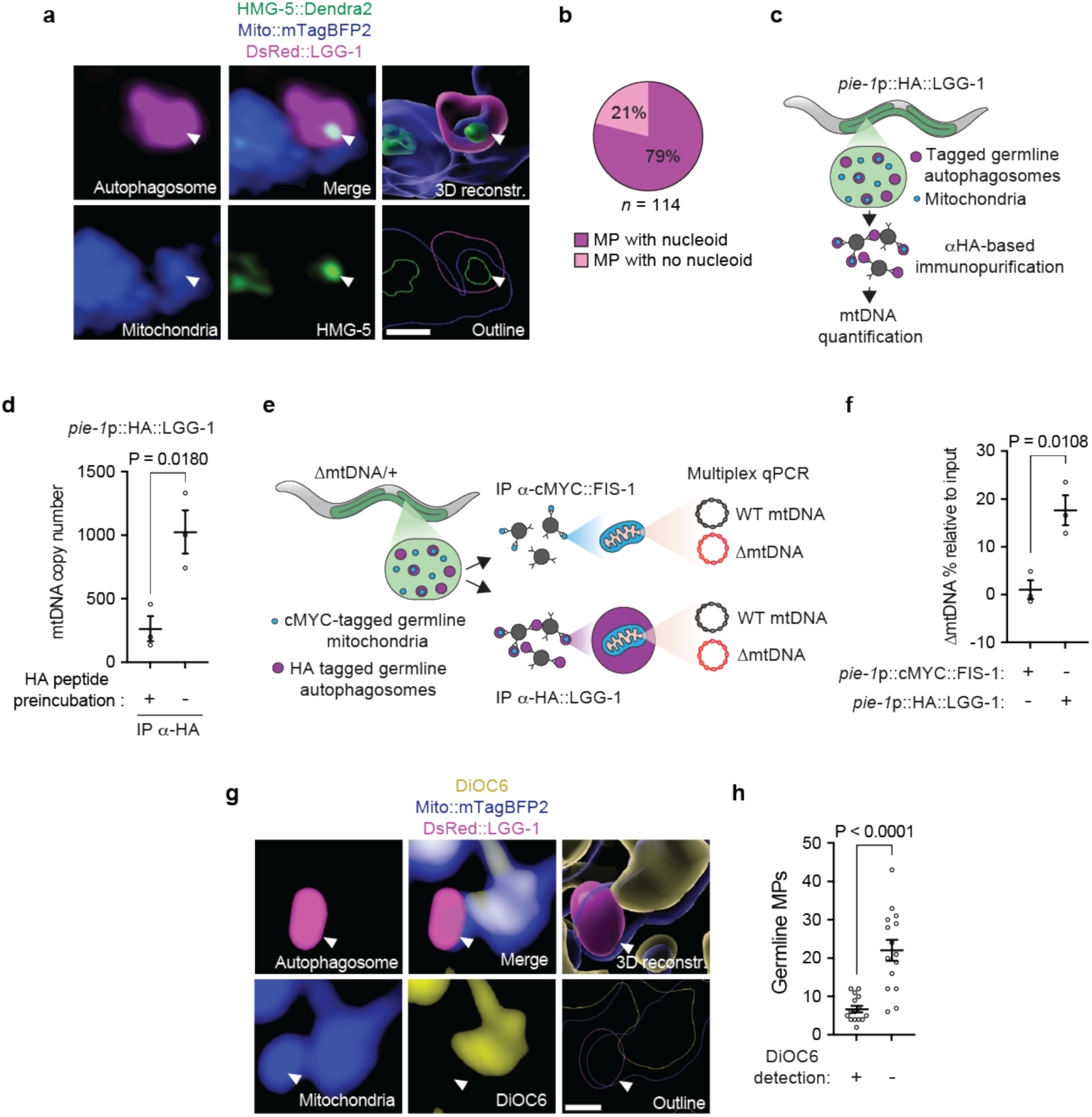
Germline Mitophagosomes Selectively Target and Engulf Mitochondria Harboring Mutant mtDNAs in the Germline. **a,** Representative image and three-dimensional reconstruction of a germline MP containing a protein–mtDNA nucleoid complex labelled by HMG-5::Dendra2 (green). Scale bar, 1 µm. **b,** Quantification of the fraction of germline MPs containing nucleoids. **c,** Schematic representation of the approach used to selectively purify germline autophagosomes. Haemagglutinin (HA)-tagged LGG-1 was specifically expressed in the germline under the *pie-1* promoter (*pie-1*p). HA-tagged LGG-1 was then immunoprecipitated from populations of animals using anti-HA antibody conjugated magnetic beads and the mtDNA contained within this fraction was analyzed by quantitative PCR (qPCR). **d,** qPCR analysis of the immunoprecipitated fractions showing enrichment of mtDNA. For the control condition, anti-HA magnetic beads were pre-incubated with HA peptide. Bars represent mean ± SEM; n = 3 independent experiments; unpaired two-tailed Student’s t-test. **e,** Schematic representation of the approach used to analyze ΔmtDNA content within germline autophagosomes. Germline-specific HA-tagged LGG-1 was expressed in animals carrying ΔmtDNA, and autophagosomes were immunoprecipitated using anti-HA magnetic beads. ΔmtDNA content in the purified fraction was quantified using multiplex qPCR. As a control, a similar approach was performed using germline-specific cMYC-tagged FIS-1 to immunopurify the total germline mitochondrial pool. **f,** Quantification of ΔmtDNA heteroplasmy content in immunoprecipitation fractions, relative to whole-animal inputs. Bars represent mean ± SEM; n = 3 independent experiments; unpaired two-tailed Student’s t-test. **g,** Representative image and three-dimensional reconstruction of germline mitochondria, autophagosomal membrane, and DiOC6 staining. Scale bar, 1 µm. **h,** Quantification of the proportion of DiOC6-negative germline MPs. Bars represent mean ± SEM; n = 3 independent experiments; unpaired two-tailed Student’s t-test.

We next asked whether these germline MPs were enriched for mutant genomes. To this end, we immunopurified autophagosomes from ΔmtDNA/+ animals and quantified heteroplasmy (the % of ΔmtDNA copies relative to wild type mtDNAs) in these fractions using a multiplex probe-based qPCR assay^26^ (Fig. 2e). Heteroplasmy levels of ΔmtDNA were significantly higher in germline MPs compared to the total pool of mitochondria in the germline that we isolated in parallel via immunopurification of a germline-specific cMYC fused to FIS-1, an outer mitochondrial membrane protein (Fig. 2f and Extended Data Fig. 2b). Thus, mutation-enriched mitochondria in the germline are preferentially sequestered by MPs.

We found that this selectivity arises from targeted engagement of defective germline mitochondria. Using the membrane potential-sensitive dye DiOC6^27^ in live animals, we observed that autophagosomal membranes preferentially assembled around germline mitochondria with reduced membrane potentials relative to neighboring organelles (Fig. 2g and h). Moreover, the mitochondrial uncoupler carbonyl cyanide 3-chlorophenylhydrazone (CCCP) induced germline MP formation (Extended Data Fig. 3a,b), together suggesting that autophagosomes engage depolarized mitochondria in the germline that harbor mutant genomes.

### Mutant mtDNA-Bearing Mitochondria Activate PINK-1/PDR-1-Dependent Mitophagy in the Germline

To define the molecular basis of germline mitophagy, we examined the canonical PINK1/Parkin pathway (PINK-1 and PDR-1 in *C. elegans*), in which stabilized PINK1 on the mitochondrial outer membrane recruits Parkin to initiate ubiquitin-dependent autophagosomal clearance^18^ (Extended Data Fig. 3c). Multiple lines of evidence indicate that defective mitochondria enriched for mutant mtDNAs directly engage this pathway in germ cells. Loss of both *pink-1* and *pdr-1*, which are normally expressed in the germline (Extended Data Fig. 3d), did not affect basal MP levels (Extended Data Fig. 3e,f), but strongly suppressed MP formation induced by either CCCP (Extended Data Fig. 3a,b) or ΔmtDNA (Fig. 3a,b), demonstrating that PINK-1/PDR-1 activity is specifically required for stress- and mutation-induced mitophagy. Furthermore, endogenous tagging of PDR-1 with mNeonGreen revealed its colocalization with germline MPs, with enhanced association in the presence of ΔmtDNA (Fig. 3c,d and Extended Data Fig. 3g,h). In addition, GFP-tagged ubiquitin was recruited to germline mitochondria during paraquat-induced mitochondrial oxidative stress in a *pdr-1*-dependent manner (Extended Data Fig. 3i,j), indicating activation of ubiquitin-mediated mitophagy.

**Fig. 3:**
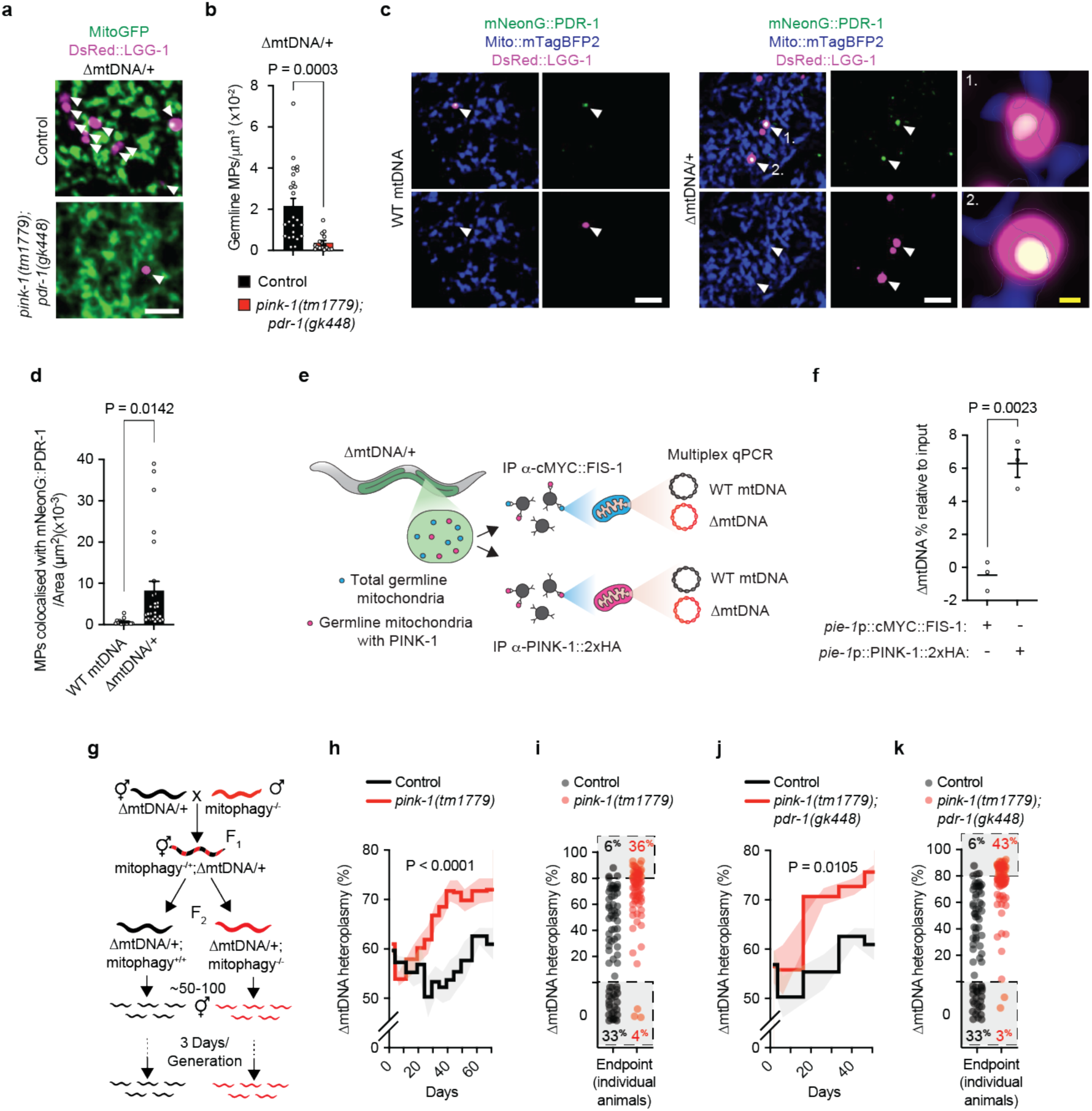
Germline Mitochondrial Harboring Mutant mtDNAs Engage the PINK-1/PDR-1 Mitophagy Pathway. **a,** Representative images of MPs in the germline of control and *pink-1(tm1779);pdr-1(gk448)* mutant animals, both carrying ΔmtDNA. MPs are indicated with white arrowheads. Scale bar, 5 µm. **b,** Quantification of germline MP abundance normalized to mitochondrial volume of the germline of control and *pink-1(tm1779);pdr-1(gk448)* animals, both carrying ΔmtDNA. Columns are mean ± SEM; n = 24 (control), n = 16 (*pink-1(tm1779);pdr-1(gk448))* germlines per condition (3 independent experiments); unpaired two-tailed Student’s t-test. **c,** Representative images of mNeonG::PDR-1 colocalizing with MPs in the germline, in the presence or absence of ΔmtDNA. Right, two representative magnified views of events 1 and 2 are shown. MPs are indicated with white arrowheads. White scale bars, 5 µm; yellow scale bar 0.5 µm. **d,** Quantification of mNeonG::PDR-1 and MP colocalization events in animals carrying wild-type mtDNA or ΔmtDNA. Columns are mean ± SEM; n = 15 (wild-type (WT) mtDNA) and 27 (ΔmtDNA) germlines (3 independent experiments); unpaired two-tailed Student’s t-test. **e,** Schematic representation of the approach used to selectively purify germline mitochondria associated with PINK-1::2xHA and analyze their ΔmtDNA content. PINK-1::2xHA was specifically expressed in the germline of animals carrying ΔmtDNA and immunoprecipitated using anti-HA magnetic beads. ΔmtDNA heteroplasmy was quantified from purified fractions using multiplex qPCR. As a control, a similar approach was performed using germline-specific cMYC-tagged FIS-1 to isolate the total germline mitochondrial pool. Bars represent mean ± SEM; n = 3 independent experiments; unpaired two-tailed Student’s t-test. **f,** Quantification of ΔmtDNA heteroplasmy content in immunoprecipitation fractions, relative to whole-animal inputs. Bars represent mean ± SEM; n = 3 independent experiments; unpaired two-tailed Student’s t-test. **g,** Schematic representation of the experimental design used to evaluate the contribution of genes that mediate mitophagy to ΔmtDNA inheritance. Crosses between ΔmtDNA/+ hermaphrodites and males genetically deficient for mitophagy genes generated F2 siblings that were either homozygous wild type (control, black) or mutant (red) for mitophagy genes. ΔmtDNA heteroplasmy was quantified across succesive generations in populations (∼10,000 animals) by transferring 50–100 animals at every new generation (every 3 days). **h,** qPCR analysis of ΔmtDNA heteroplasmy across successive generations in control and *pink-1(tm1779)* animals. Data represent the mean of ≥4 independent experiments; shaded areas indicate ± SEM. Statistical analysis was performed using two-way ANOVA. **i,** Endpoint qPCR analysis of ΔmtDNA heteroplasmy in ∼72 individual animals from control and *pink-1(tm1779)* populations. **j,** qPCR analysis of ΔmtDNA heteroplasmy across successive generations in control and *pink-1(tm1779);pdr-1(gk448)* animals. Data represent the mean of 4 independent experiments; shaded areas indicate ± SEM. Statistical analysis was performed using two-way ANOVA. **k,** Endpoint qPCR analysis of ΔmtDNA heteroplasmy in ∼72 individual animals from control and *pink-1(tm1779);pdr-1(gk448)* populations.

Finally, immunoprecipitation of PINK-1::2xHA that we expressed exclusively in the germline, revealed selective enrichment of ΔmtDNA compared to total germline mitochondria (Fig. 3e,f and Extended Data Fig. 3k), consistent with preferential stabilization of PINK-1 on mutation-enriched organelles. Together, these findings establish PINK-1/PDR-1 as the sensor-effector axis linking mtDNA mutation-associated mitochondrial dysfunction to selective mitophagy in the germline.

### Modulating PINK-1 and PDR-1 Directly Influences the Transmission of Mutant mtDNAs Across Generations

To test whether PINK-1-mediated mitophagy limits the inheritance of mutant mtDNAs across multiple generations, we tracked ΔmtDNA in lineages derived from crosses between ΔmtDNA*/+* hermaphrodites and *pink-1*-deficient males. Heteroplasmy was quantified in 13 successive generations, each consisting of a large population of animals (∼10,000) that were derived directly from F2 siblings either genetically wild type or deficient for *pink-1* (Fig. 3g). The loss of *pink-1* caused a rapid and progressive increase in ΔmtDNA heteroplasmy relative to controls (Fig. 3h), and endpoint analysis of ∼72 individuals revealed a marked shift in heteroplasmy distribution in *pink-1*-deficient lineages, with a greater fraction of animals carrying very high ΔmtDNA burdens (36% with greater than 80% heteroplasmy) and fewer animals that had cleared ΔmtDNA (4%), compared with controls (6% very high heteroplasmy and 33% cleared) (Fig. 3i). Similar heteroplasmy effects were observed in *pink-1;pdr-1* double mutants (Fig. 3j,k), demonstrating that PINK-1/PDR-1-mediated mitophagy actively restrains the intergenerational expansion of mutant mitochondrial genomes.

We next asked whether enhancing mitophagy could actively purge mutant mtDNAs from the germline, thereby reducing their transmission across generations. To test this, we generated animals that co-overexpressed *pink-1* and *pdr-1* from a single integrated MosSCI^28^ transgene driven by a ubiquitous promoter (*his-72*p), with expression monitored using a spliced nuclear-localized GFP reporter (Fig. 4a and Extended Data Fig. 4a). These animals exhibited a marked increase in germline MP abundance, consistent with elevated mitophagy activity (Fig. 4b,c). Moreover, when crossed into ΔmtDNA*/+* animals with initially very high heteroplasmy levels (∼80–90%; Fig. 4d), the enhanced mitophagy drove a rapid and progressive decline in ΔmtDNA levels across generations, culminating in complete elimination by generation nine (Fig. 4e). Notably, ΔmtDNA remained undetectable over a further ten generations, even after removal of the *pink-1/pdr-1* transgene by outcrossing (Fig. 4f), indicating complete and stable eradication from the matrilineal lineage. In contrast, control lineages lacking the *pink-1*/*pdr-1* transgene remained stably heteroplasmic at very high ΔmtDNA levels across all generations (Fig. 4e,f).

**Fig. 4:**
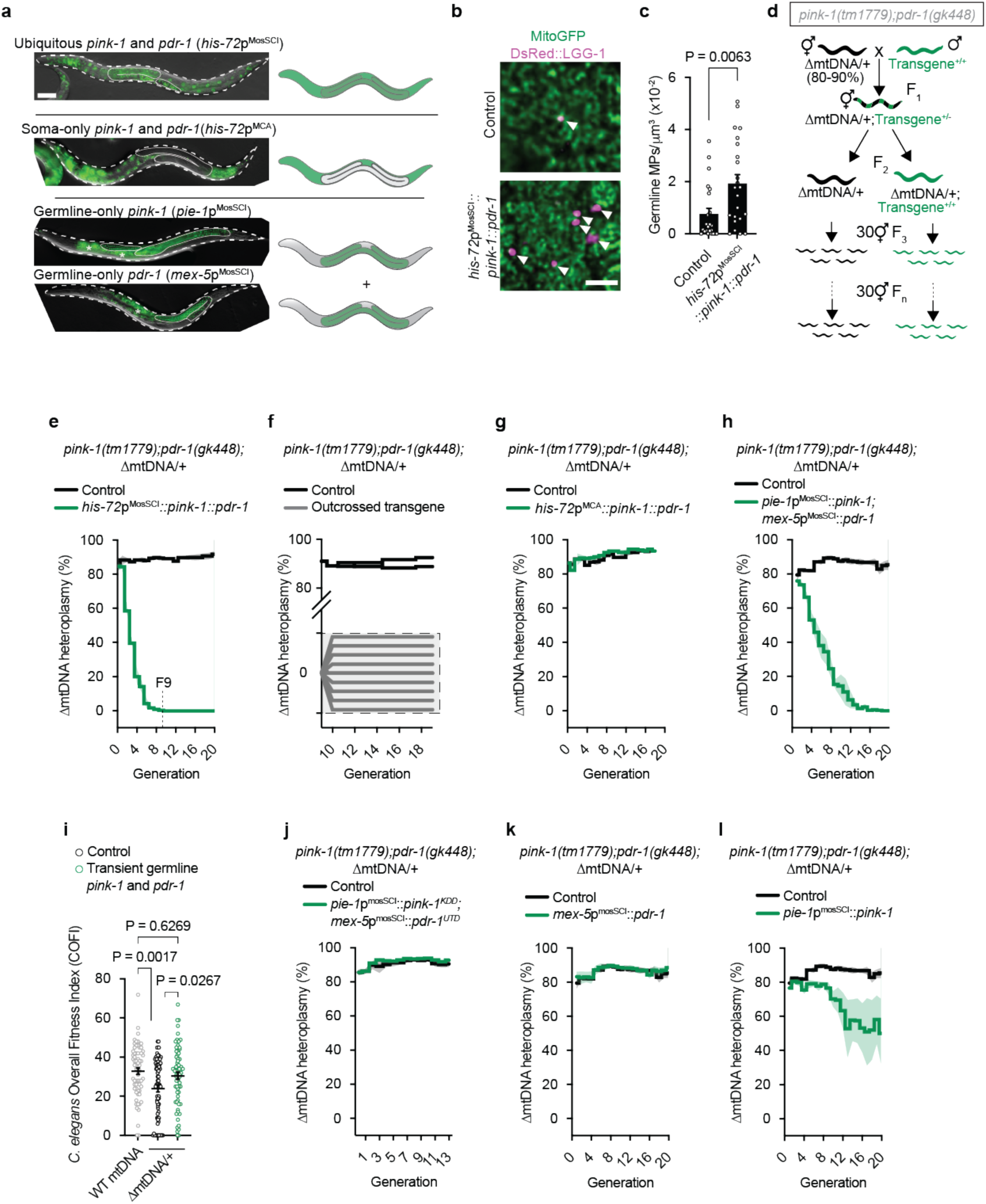
Enhancing PINK-1 and PDR-1 Germline Activities Removes Mutant mtDNA from Matrilineal Lineages. **a**, Representative images and schematic representations of transgene expression patterns in adult animals. Animals are outlined with a white dashed line, and the germline is outlined with a solid white line. Asterisks indicate autofluorescence from intestinal cells. Scale bar, 50 µm. **b,** Representative images of germline MPs in the presence or absence of *his-72*p^MosSCI^*::pink-1::pdr-1*. MPs are indicated with white arrowheads. Scale bar, 5 µm. **c,** Quantification of germline MPs normalized to mitochondrial volume in the germline in the presence or absence of *his-72*p^MosSCI^*::pink-1::pdr-1*. Columns represent mean ± SEM; n = 23 germlines per condition (3 independent experiments); unpaired two-tailed Student’s t-test. **d,** Schematic representation of the experimental design used to determine the effect of *pink-1* or *pdr-1* overexpression on ΔmtDNA inheritance. *pink-1(tm1779);pdr-1(gk448)* animals carrying high levels of ΔmtDNA (∼80–90%) were crossed with strains carrying *pink-1* and *pdr-1* transgenes generating F2 siblings that were either homozygous non-transgenic (black) or transgenic (green). ΔmtDNA heteroplasmy was quantified across successive generations by transferring 30 animals at every new generation. (**e**-**h**, **j**-**l**) qPCR analysis of ΔmtDNA heteroplasmy across successive generations in the genetic backgrounds depicted. Data represent the mean of 3 independent experiments (**e**, **g**, **h**, **j**-**l**) or 2 independent experiments for the control and 8 independent experiments for the outcrossed condition (**f**); shaded areas indicate ± SEM. **i,** Quantification of the *C. elegans* Overall Fitness Index (COFI; see Methods) in animals carrying wild-type mtDNA (WT; grey circles), control animals carrying ΔmtDNA propagated for 20 generations (black circles), and animals carrying ΔmtDNA and transiently exposed to germline overexpression of *pink-1* and *pdr-1* for 20 generations (green circles). Bars are mean ± SEM; n = 3 independent experiments; one-way ANOVA with Šídák’s multiple comparisons test.

To distinguish whether ΔmtDNA elimination was driven by enhanced mitophagy in the germline or the soma, we restricted *pink-1* and *pdr-1* overexpression to each tissue in animals lacking the endogenous genes. Somatic-only overexpression was achieved by generating animals harboring a multi-copy array (MCA) of the same transgene used above, which underwent germline silencing^29^ (Fig. 4a and Extended Data Fig. 4a), and was insufficient to reduce intergenerational ΔmtDNA loads (Fig. 4g). In contrast, germline-specific overexpression, achieved using single-copy integrated *pink-1* and *pdr-1* transgenes driven by overlapping germline promoters (*pie-1*p and *mex-5*p) (Fig. 4a and Extended Data Fig. 4a), rapidly reduced ΔmtDNA heteroplasmy across generations (Fig. 4h). Similar intergenerational reductions in heteroplasmy were observed in animals carrying the *mpt1/+* mtDNA mutation (Fig. 1f and Extended Data Fig. 4b,c), indicating that germline mitophagy can selectively purge genetically distinct mtDNA mutations. Importantly, genetically enhancing germline mitophagy did not reduce fecundity (Extended Data Fig. 4d) and completely eliminated ΔmtDNA by generation 20, restoring organismal fitness (Fig. 4i) and demonstrating that this advantage arises from selective mitochondrial quality control rather than selection at the level of whole organisms.

Consistent with the requirement of each enzyme’s mitophagy activities, disruption of either the kinase activity of transgenic PINK-1 or the ubiquitin ligase activity of transgenic PDR-1 abolished their ability to reduce intergenerational heteroplasmy (Fig. 4j and Extended Data Fig. 4e,f). Moreover, while *pink-1* and *pdr-1* co-overexpression produced robust and reproducible declines in intergenerational ΔmtDNA heteroplasmy, overexpression of *pdr-1* alone in *pink-1;pdr-1* double mutants failed to do so (Fig. 4k) and overexpression of *pink-1* alone resulted in only modest and variable intergenerational effects (Fig. 4l). As such, these findings support a model in which PINK-1/PINK1 activation is required to initiate the ubiquitin-mediated pathway, with PDR-1/Parkin amplifying PINK1-dependent phospho-ubiquitination (Extended Data Fig. 3c), and accelerating the elimination of mutant mtDNAs across generations.

Finally, germline-specific overexpression of alternative E3 ubiquitin ligases (*ari-1.1*/ARIH1, *siah-1*/SIAH1, or *wwp-1*/SMURF-1; Extended Data Fig. 4a), which are expressed in the germline (Extended Data Fig. 4g) and have been implicated in mitophagy, did not enhance PINK-1-mediated ΔmtDNA clearance (Extended Data Fig. 4h-j), indicating that a specific PINK-1/PDR-1 axis acts in the germline to facilitate intergenerational mtDNA selection.

### FIS-2 is Required for Mitophagy-Driven Intergenerational Selection Against Mutant mtDNAs

Given that PINK-1/PDR-1-dependent mitophagy selectively targets mutation-enriched mitochondria, we next asked whether mitochondrial partitioning is required for their efficient recognition and elimination. Mitochondrial fragmentation is a key determinant of mitophagy efficiency and selectivity, facilitating autophagosomal engulfment and the segregation of mitochondrial genomes into genetically resolvable units. Peripheral mitochondrial fission is particularly important in this context; in mammals, it is mediated by Fis1 and generates small mitochondrial fragments enriched for damaged material destined for mitophagy^30^.

Consistent with a heightened requirement for mitochondrial quality control, germline mitochondria were more fragmented than those in somatic tissues (Extended Data Fig. 5a-d). Moreover, animals deficient for *fis-2* (the *C. elegans* ortholog of Fis1^31,32^) exhibited increased mitochondrial branch length and a marked reduction in very small mitochondrial particles (<0.2 µm^2^) within the germline (Fig. 5a-c), consistent with defects in peripheral fission. Loss of *fis-2* also strongly suppressed ΔmtDNA-induced MP formation in the germline (Fig. 5d,e) and strongly impaired the intergenerational reduction in ΔmtDNA heteroplasmy driven by germline-specific *pink-1* and *pdr-1* overexpression (Fig. 5f). Given that mtDNA nucleoids were frequently detected within mitochondrial particles ≤0.2 µm^2^ (Extended Data Fig. 5e,f), these findings support a model in which FIS-2-dependent peripheral fission partitions mtDNA into small mitochondrial units, thereby enabling selective mitophagic elimination of mutant genomes.

**Fig. 5:**
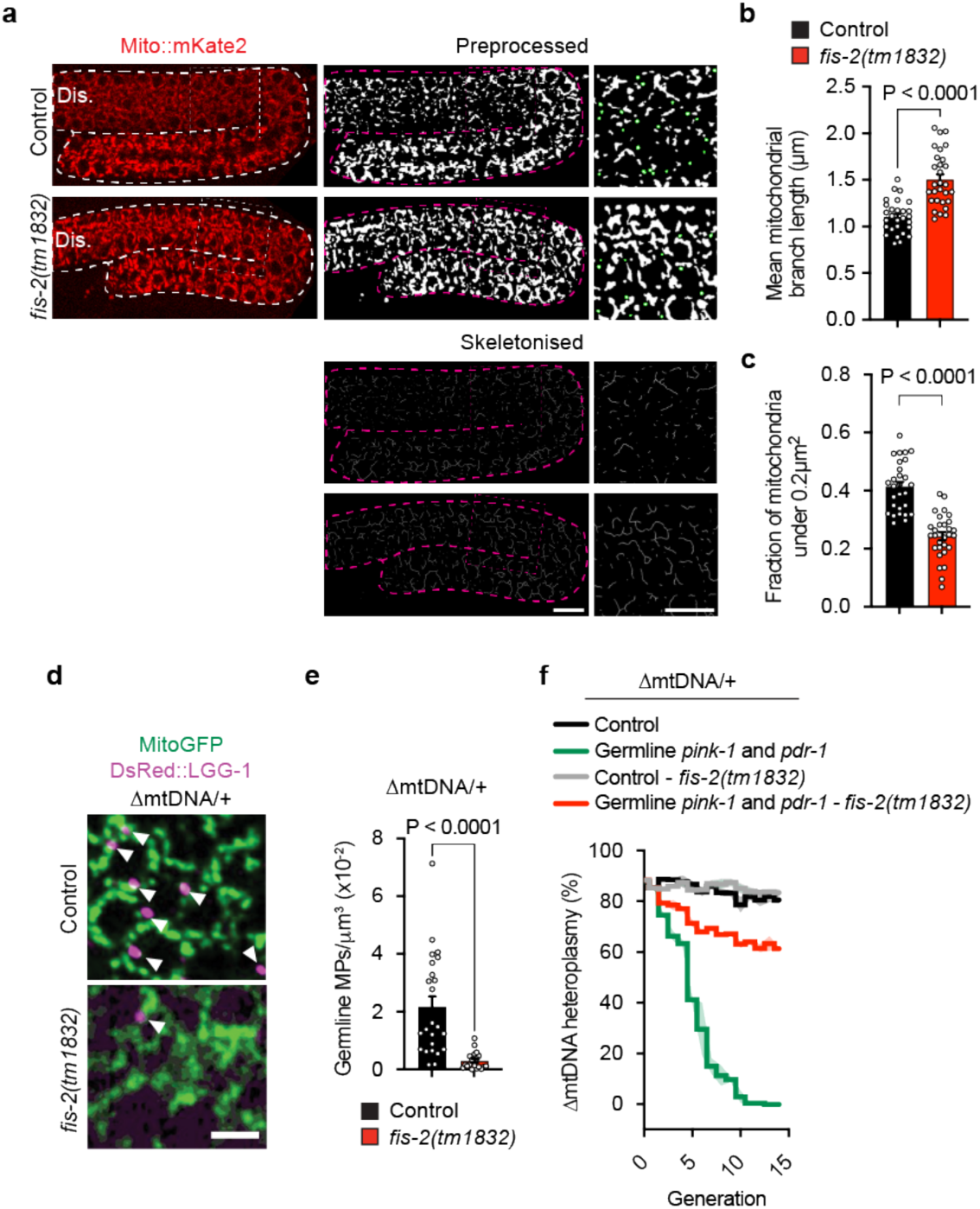
FIS-2 is Required for Mitophagy-Driven Selection Against Mutant mtDNAs in the Germline. **a,** Representative images of germline (outlined with bold dashed lines) mitochondria in control and *fis-2(tm1832)* animals. Images were preprocessed for mitochondrial size analysis and skeletonized for mitochondrial network analysis. A thin dashed square marks the zoomed region, where mitochondria <0.2 µm² are highlighted in green in the preprocessed image. Dis.: distal germline. Scale bar, 10 µm. **b,** Quantification of mean germline mitochondrial branch length in control and *fis-2(tm1832)* animals from skeletonized germline mitochondrial networks. Columns represent mean ± SEM; n = 28 (control) and 29 (*fis-2*) germlines (3 independent experiments); unpaired two-tailed Student’s t-test. **c,** Quantification of the fraction of germline mitochondria <0.2 µm² in control and *fis-2(tm1832)* animals from preprocessed images. Columns are mean ± SEM; n = 28 (control) and 29 (*fis-2*) germlines (3 independent experiments); unpaired two-tailed Student’s t-test. **d,** Representative images of germline MPs of control and *fis-2(tm1832)* animals, both carrying ΔmtDNA. MPs are indicated with white arrowheads. Scale bar, 5 µm. **e,** Quantification of germline MPs normalized to mitochondrial volume in the germline of control and *fis-2(tm1832*) animals, both carrying ΔmtDNA. Columns are mean ± SEM; n = 24 (control) and 19 (*fis-2*) germlines (3 independent experiments); unpaired two-tailed Student’s t-test. **f,** qPCR analysis of ΔmtDNA heteroplasmy across successive generations in animals carrying ΔmtDNA and germline overexpression of *pink-1* and *pdr-1*, in wild-type and *fis-2(tm1832)* backgrounds. Data represent the mean of 3 independent experiments; shaded areas indicate ± SEM.

### Receptor-Mediated Mitophagy via DCT-1 Independently Eliminates ΔmtDNA Across Generations

Despite its central role, we noticed that PINK-1/PDR-1-dependent mitophagy did not fully account for the mitophagic response to mtDNA mutations. Notably, PDR-1::mNeonGreen did not colocalize with all germline MPs (Extended Data Fig. 3g,h), and ΔmtDNA-induced MP formation was only partially suppressed by loss of both *pink-1* and *pdr-1* (Fig. 3a,b), suggesting the presence of additional mitophagy pathways. We therefore examined receptor-mediated mitophagy regulated by DCT-1, the *C. elegans* ortholog of BCL interacting protein 3 (BNIP3). Similarly to mutations in *pink-1* and *pdr-1*, loss of *dct-1* attenuated ΔmtDNA-induced MP formation (Fig. 6a,b) and increased heteroplasmy across generations, resulting in a greater proportion of animals with high ΔmtDNA burdens (Fig. 6c,d).

**Fig. 6:**
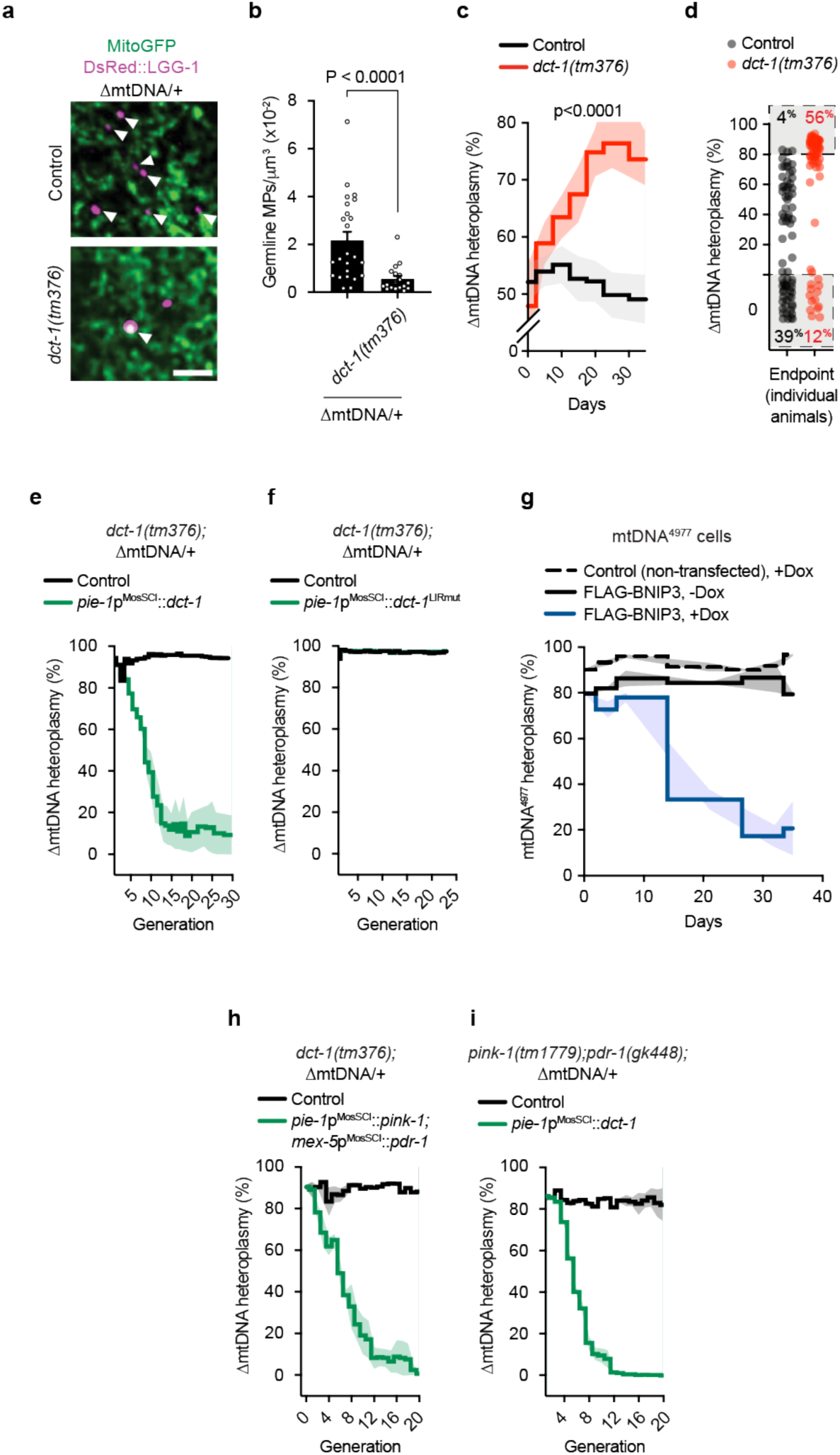
Receptor-Mediated Mitophagy via DCT-1 Eliminates ΔmtDNA Intergenerationally. **a,** Representative images of germline MPs of control and *dct-1(tm376)* animals, both carrying ΔmtDNA. MPs are indicated with white arrowheads. Scale bar, 5 µm. **b,** Quantification of germline MPs normalized to mitochondrial volume in the germline of control and *dct-1(tm376)* animals, both carrying ΔmtDNA. Columns are mean ± SEM; n = 24 (control) and 18 (*dct-1*) germlines (3 independent experiments); unpaired two-tailed Student’s t-test. **c,** qPCR analysis of ΔmtDNA heteroplasmy across successive generations in control and *dct-1(tm376)* animals. Data represent the mean of ≥4 independent experiments; shaded areas indicate ± SEM. Statistical analysis was performed using two-way ANOVA. **d,** Endpoint qPCR analysis of ΔmtDNA heteroplasmy in ∼72 individual animals from control and *dct-1(tm376)* populations. (**e**, **f**, **h**, **i**) qPCR analysis of ΔmtDNA heteroplasmy across successive generations in the genetic backgrounds depicted. Data represent the mean of 3 independent experiments; shaded areas indicate ± SEM. **g,** qPCR analysis of mtDNA^4977^ heteroplasmy across successive passages in cell lines carrying the mtDNA deletion and expressing a doxycycline-inducible FLAG-BNIP3 transgene. −Dox, no doxycycline treatment; +Dox, doxycycline treatment. Dashed line, non-transfected control +Dox; black line, FLAG-BNIP3 −Dox; blue line, FLAG-BNIP3 +Dox. Data represent the mean of three independent experiments; shaded areas indicate ± SEM.

In addition, germline-specific overexpression of *dct-1* drove rapid intergenerational elimination of ΔmtDNA (Fig. 6e) in a manner dependent on its LC3-interacting region (LIR)^33^ (Fig. 6f, Extended Data Fig. 6a), indicating that recruitment of autophagosomal membranes is required for this activity. Consistent with a conserved mechanism, BNIP3 overexpression in a cell line carrying the Kearns–Sayre syndrome (KSS)–associated mtDNA deletion (ΔmtDNA^4977^) similarly led to a progressive decline in heteroplasmy across passages (Fig. 6g and Extended Data Fig. 6b,c). Although BNIP3-mediated mitophagy is typically activated under stress conditions, such as hypoxia, where it accumulates on the mitochondrial outer membrane and promotes mitochondrial clearance^34^, the molecular basis of substrate selectivity remains incompletely understood^35^. Despite this, these findings suggest that BNIP3/DCT-1 upregulation is sufficient to drive selective elimination of mutant mtDNAs across both cell passages and generations.

Finally, although BNIP3/DCT-1 has been proposed to function within the PINK1/Parkin mitophagy pathway – either via Parkin-mediated ubiquitination that enhances GABARAP binding^36^ or by stabilizing PINK1^37,38^ – our genetic analyses indicate that these pathways operate independently in the germline. Intergenerational ΔmtDNA clearance driven by *pink-1*/*pdr-1* co-overexpression persisted in *dct-1* mutants (Fig. 6h), and conversely, *dct-1*-mediated elimination of ΔmtDNA was unaffected in *pink-1;pdr-1* double mutants (Fig. 6i). Together, these findings demonstrate that the germline deploys two genetically separable mitophagy arms – PINK-1/PDR-1- and DCT-1-mediated – that act in parallel to eliminate mutant mtDNAs, thereby reinforcing the fidelity of mitochondrial genome transmission across generations.

### Germline Mitophagy is Triggered Cell-Autonomously by mtDNA Damage

Because MP formation was induced in the germline of animals harboring mutant mtDNAs throughout all tissues^26^ (Fig. 1g,h), we asked whether germ cells directly sense mtDNA defects and trigger mitophagy in a cell-autonomous manner. To this end, we induced mtDNA double-strand breaks (mtDSBs) in a tissue-specific manner using the mitochondrial-targeted nuclease ^MTS^PstI^39^ expressed in defined cell types (Extended Data Fig. 7a). Induction of mtDSBs exclusively within neurons or body wall muscle cells failed to activate germline MPs (Extended Data Fig. 7b-e), whereas germline-restricted mtDNA damage robustly increased MP abundance (Fig. 7a,b), indicating that germ cells autonomously sense and respond to mtDNA defects.

**Fig. 7:**
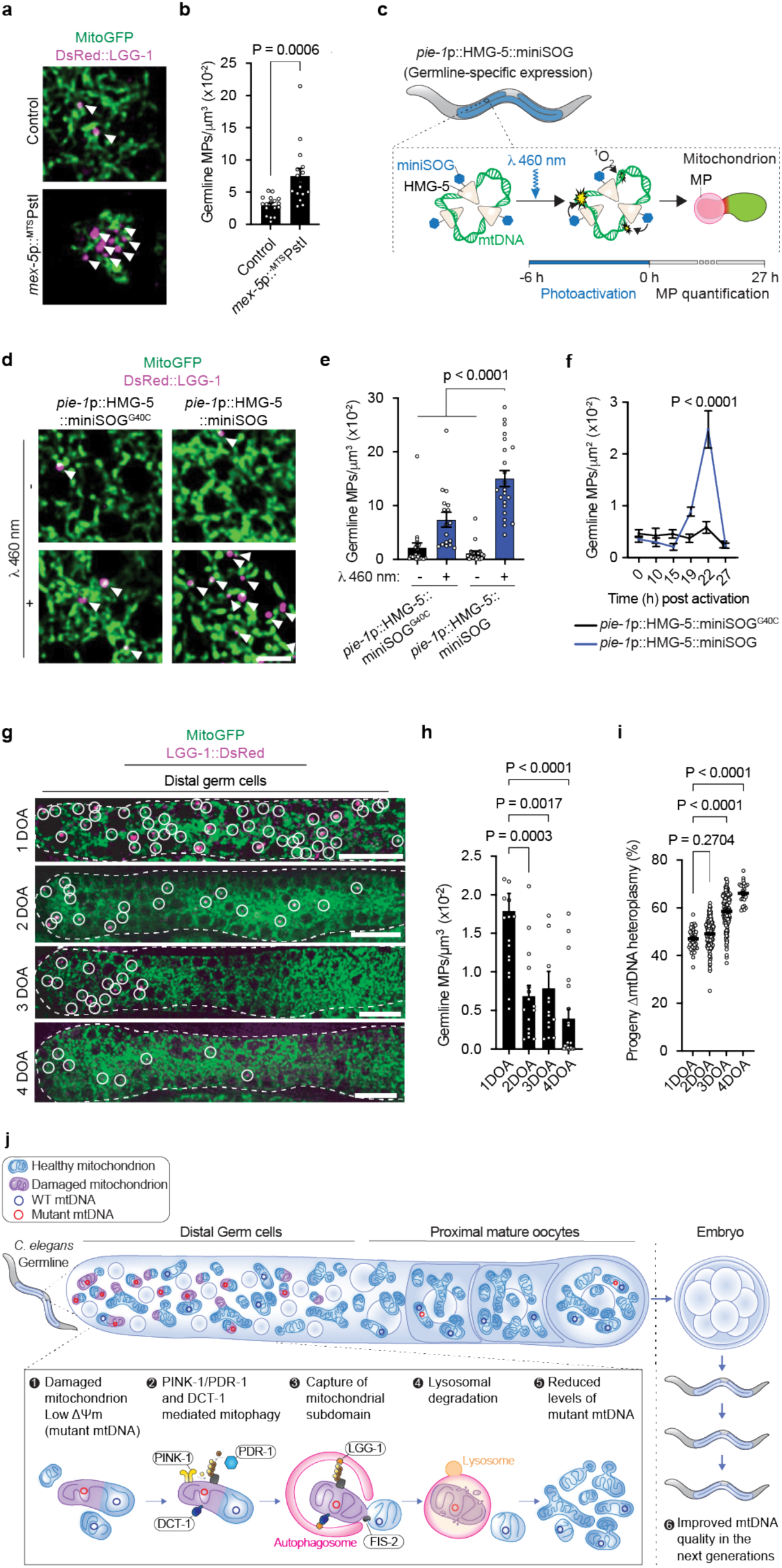
Germline Mitophagy Responds Cell-Autonomously to mtDNA Defects and Declines During Aging. **a,** Representative images of germline MPs in the presence or absence of *mex-5*p::^MTS^PstI. MPs are highlighted with white arrowheads. Scale bar, 5 µm. **b,** Quantification of germline MPs normalized to mitochondrial volume of the germline in the presence or absence of *mex-5*p::^MTS^PstI. Columns are mean ± SEM; n = 17 (control), n = 15 (^MTS^PstI) germlines per condition (3 independent experiments); unpaired two-tailed Student’s t-test. **c,** Schematic representation of optogenetic induction of germline mtDNA oxidative lesions. HMG-5 (TFAM) is fused to the mini singlet oxygen generator (miniSOG) and expressed under the germline-specific promoter *pie-1* (*pie-1*p::HMG-5::miniSOG). Upon photoactivation with 460 nm light, miniSOG locally generates ^1^O_2_, inducing oxidative mtDNA damage. **d,** Representative images of germline MPs in animals expressing *pie-1*p::HMG-5::miniSOG or the catalytically inactive variant *pie-1*p::HMG-5::miniSOG^G40C^, with or without 460 nm light photoactivation. MPs are indicated with white arrowheads. Scale bar, 5 µm. **e,** Quantification of germline MPs normalized to mitochondrial volume of the germline in animals expressing *pie-1*p::HMG-5::miniSOG or the catalytically inactive variant *pie-1*p::HMG-5::miniSOG^G40C^, with or without 460 nm light photoactivation. Columns are mean ± SEM; n = 21 (miniSOG^G40C^, non-illuminated), 17 (miniSOG^G40C^, illuminated), 20 (miniSOG, non-illuminated), and 22 (miniSOG, illuminated) germlines (2 independent experiments); one-way ANOVA with Šídák’s multiple comparisons test. **f,** Quantification of MPs normalised to germline area (µm^2^) across successive timepoints following photoactivation of HMG-5::miniSOG (blue line) or the catalytically inactive HMG-5::miniSOG^G40C^ control (black line). Data are mean ± SEM; HMG-5::miniSOG: *n* = 13 (0 h), 11 (10 h), 11 (15 h), 23 (19 h), 19 (22 h), and 24 (27 h) germlines. HMG-5::miniSOG^G40C^: *n* = 12 (0 h), 9 (10 h), 11 (15 h), 26 (19 h), 18 (22 h), and 22 (27 h) germlines; one-way ANOVA with Sidak’s multiple comparisons test. **g,** Representative images showing the evolution of MP abundance in the distal germline region during aging, from 1-day-old adult (1DOA) to 4-day-old adult (4DOA). MPs are indicated with white circles. Scale bar, 20 µm. **h,** Quantification of germline MPs normalized to mitochondrial volume in the germline of animals during aging. Columns represent mean ± SEM; n = 15 (1DOA), 16 (2DOA), 13 (3DOA), and 20 (4DOA) germlines (3 independent experiments); one-way ANOVA with Šídák’s multiple comparisons test. **i,** Quantification of ΔmtDNA heteroplasmy in progeny produced at different time points during aging. Bars represent mean ± SEM; n = 42 (1DOA), 129 (2DOA), 120 (3DOA), and 25 (4DOA) progenies (1 of 3 independent experiments is shown here); one-way ANOVA with Šídák’s multiple comparisons test. **j,** Proposed model of germline mitophagy selectively eliminating mutant mtDNA within the germline, thereby restricting their supply to unfertilized oocytes and transmission to the next generation.

To further examine the germline-autonomous surveillance properties of mitophagy and determine whether it responds to additional forms of mtDNA damage, we examined the effects of oxidative lesions. Treatment with paraquat, which increased mtDNA oxidative damage (Extended Data Fig. 7f) and induced *pdr-1*-dependent mitochondrial ubiquitination (Extended Data Fig. 3i,j), also promoted germline MP formation (Extended Data Fig. 7g,h). To selectively restrict oxidative damage to germline mtDNA, we developed an optogenetic approach in which the photosensitizer miniSOG was fused to HMG-5 and expressed specifically in germ cells (Fig. 7c). Upon illumination with 460-nm light, miniSOG generates singlet oxygen (^1^O_2_) within a radius comparable to that of mtDNA nucleoids (∼100 nm)^40,41^. Photoactivation of HMG-5::miniSOG induced mtDNA damage (Extended Data Fig. 7i) and robustly increased germline MP abundance relative to non-illuminated controls or animals expressing a catalytically inactive HMG-5::miniSOG^G40C^ variant (Fig. 7d,e). The inactive construct produced only a modest, non-significant increase in MPs, likely reflecting phototoxic effects of blue light or residual miniSOG activity (Extended Data Fig. 7i).

By quantifying MP formation across multiple timepoints following HMG-5::miniSOG photoactivation (Fig. 7c), we found that MPs began to accumulate within the germline between 15 and 19 hours after mtDNA damage induction (Fig. 7f), defining a temporal sequence of events in which oxidative mtDNA damage gives rise to mitochondrial dysfunction sufficient to trigger mitophagy. Together, these findings establish germline mitophagy as a cell-autonomous surveillance mechanism that senses and responds to mtDNA defects as they arise within germ cells.

### An Age-Dependent Distal Germline Mitophagy Checkpoint Controls mtDNA Inheritance

Having established a germline-autonomous surveillance role for mitophagy that selectively eliminates deleterious mitochondrial genomes and restricts their intergenerational transmission, we next asked how this process is spatially and temporally organised across the reproductive axis. In *C. elegans*, germ cells are organized along a distal-proximal gradient, with proliferating GSCs positioned distally and mature oocytes proximally (Fig. 1a). In young adults, MPs were enriched in the distal germline and further increased in response to ΔmtDNA and mitochondrial stress (Fig. 1b and Extended Data Fig. 7j), identifying this region as a focal site of mitophagy.

In support, gonad dissection revealed that distal germ cells exhibited significantly higher ΔmtDNA heteroplasmy than unfertilized oocytes from sperm-deficient *spe-26(it112)* animals^42^ (Extended Data Fig. 7k), excluding contributions from oocyte-zygote transition^22^ or embryogenesis. In contrast, *pink-1;pdr-1;dct-1* triple mutants showed elevated ΔmtDNA levels in oocytes relative to distal germ cells (Extended Data Fig. 7l), identifying the distal germline as a spatially defined mitophagy checkpoint that restricts mutant mtDNAs from being supplied to oocytes prior to fertilization. To directly test this, we restricted *pink-1* transgenic expression to the distal germline by replacing its *tbb-2* 3’ UTR with that of *fbf-1*, which confines expression to mitotically and meiotically dividing germ cells^43,44^ (Extended Data Fig. 7m). In combination with *pdr-1* overexpression – which alone does not influence heteroplasmy (Fig. 4k) – distal-restricted *pink-1* expression was sufficient to drive intergenerational reductions in heteroplasmy (Extended Data Fig. 7n).

Finally, if germline mitophagy functions as a checkpoint that filters mtDNA mutations prior to oocyte fertilization, its effectiveness may depend on how it is regulated across the reproductive window. We therefore asked whether mitophagy activity in the germline changes with age, influencing mtDNA transmission to offspring. We found that the abundance of ΔmtDNA-induced MPs in the distal germline declined markedly between days 1 and 4 of adulthood (Fig. 7g,h), spanning the reproductive period of hermaphrodites. To determine whether this decline impacts inheritance, we quantified ΔmtDNA heteroplasmy in progeny produced on successive reproductive days. Offspring derived from later broods consistently carried higher ΔmtDNA loads than those produced earlier in adulthood (Fig. 7i), indicating that age-associated attenuation of distal germline mitophagy is associated with increased transmission of mutant mitochondrial genomes.

## Discussion

Faithful transmission of genetic information through the immortal germline is fundamental to the survival of all species. Here, we show that canonical mitophagy functions as a germline surveillance mechanism that actively restricts the expansion and intergenerational transmission of deleterious mitochondrial genomes. Two mechanistically distinct pathways – PINK-1/PDR-1 (PINK1/Parkin)-mediated mitophagy and receptor-mediated mitophagy driven by the LC3-interacting receptor DCT-1 (BNIP3) – operate in parallel within GSCs to sense and respond to mtDNA mutations and damage, triggering the selective autophagosomal engulfment and removal of mitochondria enriched for mutant genomes before their incorporation into unfertilized oocytes and propagation along the matrilineal lineage. Together, these findings establish germline canonical mitophagy as a major layer of intergenerational mtDNA selection.

Our results support a model (Fig. 7j) in which germline mitophagy is driven by mitochondrial compartmentalization followed by selective elimination of mutant genomes. In this model, the peripheral fission factor FIS-2 generates genetically resolvable mitochondrial units or subdomains that independently engage PINK-1/PDR-1 signaling or DCT-1 activity to recruit LGG-1 positive autophagosomal membranes. MP formation preferentially occurs on mitochondria with reduced membrane potential and elevated mutant mtDNA loads, culminating in their delivery to acidified compartments that degrade mitochondrial contents and thereby eliminate mutant genomes. Consistent with this, enhancing PINK-1/PDR-1 or DCT-1 activity in the germline promotes the intergenerational removal of mutant genomes.

Although purifying selection of mtDNA is a conserved feature of metazoan reproduction, the existence of mechanisms that actively and selectively eliminate mutant genomes to prevent their transmission across generations remains unclear. Mitophagy in primordial germ cells (PGCs)^19^ and during the oocyte-to-zygote transition (OZT)^22^, is developmentally timed rather than mutation-responsive^19,22^, as is a similarly programmed mitophagy phase in *Drosophila* ovaries^21^. By contrast, the germline mitophagy described here is directly triggered by mtDNA mutations, oxidative damage, and mtDNA double-strand breaks. Whereas PINK-1, PDR-1, and DCT-1 do not influence basal germline mitophagy levels, they are required for mitophagy triggered by mtDNA mutations. Thus, these pathways function as a responsive surveillance mechanism that detects mitochondrial dysfunction and promotes the selective removal of mitochondria harbouring mutant mtDNAs from the heritable pool, rather than as a temporally programmed organelle remodelling event. In line with these distinct roles, developmentally programmed mitophagy is unlikely to selectively eliminate mutant genomes and is therefore unlikely to substantially influence intergenerational mtDNA inheritance in viable adult animals.

This distinction likely reflects the cellular contexts in which these processes operate. The distal germline contains proliferating GSCs characterized by active mitochondrial biogenesis and rapid mtDNA replication^19,45^, conditions that increase the likelihood of mutation and necessitate continuous mitochondrial quality surveillance. Consistent with this, germline mitophagy is mechanistically distinct from mitochondrial turnover in PGCs and during the OZT. In the PGCs, mitochondrial turnover does not involve DCT-1 and, although it requires PINK-1, proceeds independently of both PDR-1 and autophagy^19^. By contrast, OZT mitophagy occurs independently of PINK-1, PDR-1, and DCT-1 and instead depends on FUNDC-1^22^, which mediates receptor-driven mitophagy, although whether this process selectively targets mitochondria harbouring mutant mtDNAs remains unknown. Because we and others have shown that PINK-1, PDR-1/Parkin^46^ and DCT-1/BNIP3 do selectively target such organelles, these mechanistic differences raise the possibility that distinct mitophagy programmes are specialised for either selective genome surveillance or broader bulk mitochondrial turnover.

Notably, multiple models of mtDNA purifying selection invoke mitophagy-associated proteins acting through non-canonical mechanisms. For example, in *Drosophila* ovaries, PINK1 accumulation on defective mitochondria suppresses local mitochondrial protein translation, including that of PolG1, thereby limiting replication of mutant genomes^10,11^. Similarly, BNIP3 and Atg1 have been proposed to influence mtDNA selection by promoting replication of bioenergetically competent mitochondria rather than selectively degrading mutant organelles^12^. These observations suggest that mtDNA selection can occur independently of mitophagic degradation. Whether canonical mitophagy represents a broadly conserved mechanism for intergenerational mitochondrial genome surveillance therefore remains an open question. However, recent studies in mice indicate that receptor-mediated mitophagy via BCL2L13^20^ and ubiquitin-mediated mitophagy regulated by the deubiquitinase USP30 can influence mtDNA selection during maternal-zygote transition^23^. Thus, it is probable that mtDNA quality control is distributed across distinct mechanisms along the germ cell–oocyte–embryo developmental continuum. In this framework, sequential and partially overlapping mechanisms progressively would reduce the burden of deleterious mtDNAs before embryogenesis.

Despite these safeguards, mtDNA mutations persist and are transmitted across generations. Strikingly, we find that genetically enhancing germline mitophagy is sufficient to completely eliminate mutant mtDNAs over successive generations without reducing fecundity or producing overt developmental costs. This raises the question of why purifying selection is not naturally engaged at higher levels. One possibility is that excessive mitochondrial turnover imposes subtle metabolic or developmental trade-offs, necessitating dynamic tuning of mitophagy to balance mitochondrial quality control with the energetic and biosynthetic demands of germ cell proliferation and oocyte production. Indeed, we find that germline mitophagy declines with age, indicating that mitochondrial surveillance is not constitutively maximized but instead varies across the reproductive lifespan. This decline coincided with increased transmission of mutant genomes to offspring, consistent with reports that mtDNA mutation burden in human offspring increases with maternal age^6^.

Collectively, our findings establish PINK-1/PDR-1- and DCT-mediated germline mitophagy as an active surveillance mechanism that selectively removes mitochondria harboring mutant mtDNAs from the heritable mitochondrial pool. As such, it represents a fundamental determinant of mitochondrial genome inheritance, with broad implications for maternal aging, mitochondrial disease transmission, and the long-term evolutionary stability of the mitochondrial genome.

## Acknowledgements

This work was supported by an ARC Discovery Project DP200101630, ARC Future Fellowship, and Stafford Fox Senior Research Fellowship (to S.Z.), NHMRC grants GNT2047527 and GNT2048665 (to S.Z.), and an NHMRC Centre of Research Excellence in Mechanisms In NeuroDegeneration– Alzheimer’s Disease (MIND-AD CRE) GNT2035494 (to S.Z.).

## Materials and Methods

### *C. elegans* strains and maintenance

All *C. elegans* strains used, their genotype and origin are listed in Supplementary Table 1. Animals were maintained following standard methods^47^ on Nematode Growth Medium (NGM) agar plates seeded with OP50 *E. coli*.

### Transgenic strains

Mos-mediated single copy insertion (MosSCI, *foxSi* alleles) transgenic animals were produced by microinjecting 50 ng/μl of assembled MosSCI targeting plasmid in universal MosSCI strains, as previously described^28^. Prior to injection, plasmids were purified using the PureLink™ HiPure Plasmid Miniprep Kit (Invitrogen).

Extrachromosomal arrays were introduced by injection of 5 ng/μl plasmid DNA purified using the PureLink™ Quick Plasmid Miniprep Kit (Invitrogen) into the gonad of wild-type (N2) animals using standard microinjection procedures. *odr-1p::DsRed* (10 ng/μl) was used as a co-injection marker. The *foxEx104* extrachromosomal array was integrated into the *C. elegans* genome using UV irradiation following standard procedure to create *foxIs1*.

An endogenous 3xFLAG-auxin-inducible degron (AID) fused to mNeonG (mNeonGreen) was inserted in frame at the N terminus of the *pdr-1* gene using CRISPR/Cas9-mediated homology-directed repair (*fox60[3xFLAG::mNeonG::AID::pdr-1]III*). A Dendra::FLAG fusion protein tag was inserted in frame at the C terminus of the *hmg-5* gene using CRISPR-Cas9-mediated homology-directed repair (*hmg-5(fox43[hmg-5::Dendra::FLAG])IV*). The repair templates encompassed silent mutations to disrupt the PAM site and prevent Cas9 re-cleavage. Correct integration was verified by PCR and Sanger sequencing.

*foxSi277* (*pie-1p::pink-1::2xHA [wbmIs60 III]*) was generated using the SKI LODGE CRISPR/Cas9 targeted single-copy insertion strategy^48^. *pie-1p::SL2::pink-1::2xHA* was amplified from a gBlocks™ Gene Fragment (Integrated DNA Technologies, IDT) with AA395 and AA396. This DNA fragment was purified using AMPure XP beads (Beckman Coulter) and injected into WBM1119, *wbmIs60 (pie-1p::3xFLAG::dpy-10 crRNA::unc-54 3’UTR)*. The 3xFLAG tag was excluded during the recombination process.

### Constructs

The plasmids generated for this study are listed in Supplementary Table 2 and the oligonucleotide sequences in Supplementary Table 3. Oligonucleotides were supplied by IDT. Phusion™ High-Fidelity DNA Polymerase (Thermo Scientific™), T4 polynucleotide kinase (New England Biolabs, NEB), T4 DNA ligase (NEB) and the restriction enzyme *Dpn*I (NEB) were used for PCR-based cloning. For Gibson assembly^49^, Phusion DNA polymerase (NEB), Taq DNA ligase (NEB), and T5 exonuclease (NEB) were used.

Wild-type (N2) animals were lysed for 1 h at 65°C, in Mitochondrial Lysis Buffer (MLB)^50^, supplemented with 0.1 mg/ml proteinase K. genomic DNA (gDNA) was extracted with phenol:chloroform:isoamyl alcohol (25:24:1, v/v/v) following standard procedure, and used to clone *pink-1*, *pdr-1, dct-1, hmg-5* and *ari1.1* genomic sequences. The coding sequences of *pink-1*, *pdr-1*, *siah-1* and *wwp-1* were cloned from a complementary DNA (cDNA) library prepared from wild-type (N2) animals.

pSZ45 (*rgef-1p::^MTS^PstI::mKate2*) is a derivative of pSZ6 (*myo-3p::^MTS^PstI::mKate2*)^39^, in which the *myo-3* promoter was replaced with the *rgef-1* promoter amplified using primers AT22 and AT23. pSZ3 (*mex-5p::^MTS^PstI::gSL2::his-58::GFP::tbb-2 3′ UTR*) is a derivative of pSZ164 (*mex-5p::pdr-1::gSL2::his-58::GFP::tbb-2 3′ UTR*), in which *pdr-1* was replaced with *^MTS^PstI* amplified using primers AT19 and AT5.

pSZ130 *(pie-1p::pink-1::rSL2::pdr-1::gSL2::his-58::GFP::tbb-2 3’ UTR)* was created by amplifying the *pink-1* sequence from N2 gDNA using primers AA140 and AA141, the *rSL2* sequence using primers AA142 and AA143, and the *pdr-1* sequence using primers AA144 and AA145. The plasmid backbone (pSZ114) was amplified using primers AA139 and AA146, and all linear DNA fragments were joined together following the standard Gibson assembly^49^ protocol. pSZ131 *(his-72p::pink-1::rSL2::pdr-1::gSL2::his-58::GFP::tbb-2 3’ UTR)* was created by amplifying the *his-72* promoter sequence from N2 gDNA using primers AA152 and AA153. The plasmid backbone (pSZ130) was linearized using primers AA11 and AA154. The two linear DNA fragments were assembled using the standard Gibson assembly^49^ protocol. pSZ163 *(pie-1p::pink-1::gSL2::his-58::GFP::tbb-2 3’ UTR)* was created by performing a reverse PCR using pSZ130 as a template with primers AA141 and AA146. The *pie-1* promoter sequence [ACCTTTAAATAAAATCGAGA… AATCAAATTTTCTTTTCCAG] in pSZ130, was swapped with the *mex-5* promoter sequence [ATATCAGTTTTTAAAAAATT… TTGAATGTTTCAGACAGAGA], and the resulting template was used to create pSZ164 *(mex-5p::pdr-1::gSL2::his-58::GFP::tbb-2 3’ UTR)* by removing *pink-1::rSL2* using reverse PCR with the primers AA144 and AA170. pSZ124 *(mex-5p::hmg-5::miniSOG::tbb-2 3’ UTR)* was generated by MEGAWHOP^51^ cloning. The *hmg-5* sequence was amplified from gDNA using primers ABG1 and ABG2 to create a megaprimer. The megaprimer was subsequently used to replace the *his-72* sequence in pCZ886^52^ with primers ABG1 and ABG3. pSZ136 *(mex-5p::hmg-5::miniSOG^G40C^::tbb-2 3’ UTR)* was created by inserting a loss-of-function point mutation by site-directed mutagenesis, using primers abg7 and abg8. pSZ325 *(mex-5p::ari-1.1::gSL2::his-58::GFP::tbb-2 3’ UTR),* pSZ327 *(mex-5p::siah-1::gSL2::his-58::GFP::tbb-2 3’ UTR)* and pSZ328 *(mex-5p::wwp-1::gSL2::his-58::GFP::tbb-2 3’ UTR)* were created by amplifying the *ari-1.1* gDNA sequence with primers AA198 and AA199, the *siah-1* cDNA sequence with primers AA202 and AA203 and the *wwp-1* cDNA sequence with primers AA204 and AA205. PCR fragments were digested with SpeI and NotI and ligated into pSZ164, which was previously amplified with primers AA196 and AA197 and digested with NheI and NotI. pSZ200 *(pie-1p::pink-1^KDD^::gSL2::his-58::GFP::tbb-2 3’ UTR)* was created by inserting loss-of-function point mutations into *pink-1* cDNA, previously amplified with AA165 and AA129 and subcloned in pJET1.2 (CloneJET™), through two successive reverse PCRs using the primer sets AA246 and AA247 and AA248 and AA249. Then, *pink-1^KDD^* was inserted into pSZ163 by MEGAWHOP.

pSZ201 *(mex-5p::pdr-1^UTD^::gSL2::his-58::GFP::tbb-2 3’ UTR)* was generated by inserting a loss-of-function point mutation into *pdr-1* cDNA, previously amplified with primers AA168 and AA171 and subcloned in pJET1.2 (CloneJET™), through two successive reverse PCRs using primer set AA250 and AA251. *pdr-1^UTD^* was then inserted into pSZ164 by MEGAWHOP. pSZ262 *(pie-1p::dct-1::GFP::FLAG::tbb-2 3’ UTR)* was made by amplifying *dct-1* from *gDNA* with primers AA264 and AA265. The plasmid backbone (pSZ163) was amplified using primers AA269 and AA268 and the two linear DNA fragments were assembled using standard Gibson assembly^49^. The FLAG tag was then inserted by reverse PCR using primers AA271 and AA272. pSZ315 *(pie-1p::pink-1::F2A::his-58::GFP::fbf-1 3’ UTR)* was created by amplifying the *fbf-1 3’ UTR* from gDNA with primers AA306 and AA307. The *fbf-1 3’ UTR* was then inserted into pSZ163 by MEGAWHOP. pSZ352 *(pie-1p::dct-1^LDD^::GFP::FLAG::tbb-2 3’ UTR)* was generated by reverse PCR using pSZ262 as a template with primers AA347 and AA348. pSZ387 *(pie-1p::mts::mTagBFP2::tbb-2 3’ UTR)* was generated by amplifying *mTagBFP2* from pTD75^53^ with primers AA359 and AA360. The plasmid backbone (pSZ28) was amplified using the primers at23 and AA1, and the two linear DNA fragments were assembled using standard Gibson assembly^49^. pSZ38 (*pie-1p::DsRed::lgg-1::tbb-2 3’ UTR*) was created by amplifying *DsRed::lgg-1* from pNT807, a gift from N. Tavernarakis (IMBB-FoRTH, Heraklion, Greece), with primers AA59 and AA60. *DsRed::lgg-1* was then phosphorylated and blunt-end ligated into the pSZ28 backbone. pSZ437 *(pie-1p::HA::lgg-1::SL2::tomm-20::cMYC::tbb-2 3’ UTR)* was generated from a gBlocks™ Gene Fragment (Integrated DNA Technologies) containing *HA::lgg-1::SL2::tomm-20::cMYC*, which was subcloned into pJET1.2. *HA::lgg-1::SL2::tomm-20::cMYC* was subsequently amplified and inserted into the pSZ28 backbone. pSZ439 *(pie-1p::tomm-20::cMYC::tbb-2 3’ UTR)* was generated by amplifying tomm-20::cMYC from a gBlocks™ Gene Fragment with primers AA9 and AA4 and then inserting it into the pSZ28 backbone. pSZ441 *(pie-1p::HA::fis-2::tbb-2 3’ UTR)* was generated by amplifying *HA::fis-2* from a gBlocks™ Gene Fragment with primers AA37 and AA370 and then inserting it into pSZ28 backbone. pSZ442 *(pie-1p::cMYC::fis-1::tbb-2 3’ UTR)* was generated by amplifying *cMYC::fis-1* from a gBlocks™ Gene Fragment with primers AA415 and AA416 and then inserting it into the pSZ28 backbone.

### Quantification of *uaDf5* heteroplasmy

Absolute copy numbers of wild-type and *uaDf5* mtDNA (ΔmtDNA) were determined using a probe-based qPCR approach with a Rotor-Gene Q instrument with Rotor-Gene Q Pure Detection software (version 2.3.1, QIAGEN). Primers AA123, AA125, AA126 and probes AA127 HEX/ZEN/IABKFQ (Iowa Black FQ) and AA124 6-FAM/ZEN/IABKFQ were used to quantify the wild-type mtDNA and ΔmtDNA levels. The detection system was calibrated using a plasmid carrying a wild-type fragment of mtDNA (pSZ116) and an equivalent plasmid carrying the *uaDf5* deletion (pSZ66), as previously performed^26^. The cycle conditions were 95 °C for 3 min, followed by 40 cycles of two-step cycling consisting of 94 °C for 15 s and 58 °C for 40 s.

### Quantification of *mpt1* heteroplasmy

A semi-quantitative PCR approach, performed on standard animal crude lysates using the primers BG27 and BG28, was used to quantify the heteroplasmy level of *mpt1*, as previously described^54^.

### Microscopy and fluorescence intensity quantification

For fluorescence visualization and analyses, *C. elegans* were placed on a 2% agarose pad on a glass slide and immobilized in 20 µM levamisole (Sigma) with 0.1% tricaine, under a coverslip. Animals were imaged using either a Zeiss Z2 imager microscope, at 40× magnification, equipped with a Zeiss Axiocam 506 mono camera, a Yokogawa spinning disk confocal microscope at 63× magnification, via a Hamamatsu ORCA-Flash4.0 V2 sCMOS camera (photon conversion factor 0.46), or a Zeiss LSM 710 confocal microscope, at 100× magnification, with high-sensitivity BiG (GaAsP) detectors.

Confocal images were deconvolved with Huygens Professional v18.04 (Scientific Volume Imaging, http://svi.nl) run on a GPU-accelerated computer (3× NVIDIA® Tesla® V100), using the CMLE algorithm, with SNR:20 and 40 iterations. Imaris (version 10.1.1, Bitplane) was used to generate three-dimensional renderings of fluorescent images and to visualize, analyse, and quantify data. Alternatively, ImageJ^55^ (version 1.53f51) was used to quantify the fluorescence intensity of some data. The authors gratefully acknowledge the Queensland Brain Institute advanced microscopy facility for their support and assistance in this work.

### Animal treatment before imaging

Before imaging, L4 animals were exposed to 8 mM of paraquat (Sigma, 75365-73-0), 15 µM of CCCP (Sigma, 555-60-2), or DMSO (control) for 2 days at 20°C, on plates where OP50 *E. coli* bacteria were killed by UV pre-treatment as described previously ^56^. L4 animals expressing *mex-5p::*^MTS^PstI were placed at 25°C for 24 h prior to imaging in order to increase ^MTS^PstI activity. 1-day-old adult animals expressing *pie-1p::hmg-5::miniSOG* or *hmg-5::miniSOG^G40C^*, were illuminated with blue light (λ ≈ 460 nm) for 6 h, and allowed to recover for 22 h at 20°C before imaging.

### Mitochondrial GFP- ubiquitin quantification

Channel signals were thresholded using identical image settings to generate binary masks, which were combined using the ImageJ Image Calculator (AND function) to reveal colocalization events. Colocalization was quantified as total area (µm²) using particle analysis and normalized to germline area (µm²), generating a relative arbitrary unit.

### Mitophagosome quantification

Mitophagosomes in the germline were identified by the colocalization of the autophagosome marker *foxSi30* (*pie-1p::DsRed::lgg-1::tbb-2 3’ UTR (oxti444)III*) and a mitochondrial marker *foxSi6* (*pie-1p::tomm-20(mts)::GFP::tbb-2 3’ UTR (oxti179)II*). The germlines of live animals were imaged using a Yokogawa spinning disk confocal microscope at 63× magnification. Red (autophagosome) and green (mitochondria) channels were deconvolved and 3D rendered in batch using consistent threshold parameters across datasets; minor adjustments were applied when necessary to account for variability in signal intensity due to *in vivo* imaging conditions. Colocalization events were defined based on 3D object proximity. Mitochondria (green channel) and autophagosomes (red channel) were independently rendered as 3D objects. A colocalization event was scored when the surface distance between red and green objects was equal to 0 µm (contact) or when objects exhibited partial or complete volumetric overlap. Colocalization events were quantified per germline mitochondrial volume using the Object Statistics function in Imaris. Mitophagosome spatial distribution, from the distal tip to the proximal area of the germline, was manually scored using ImageJ and displayed as heat-map with GraphPad Prism (V10.2.0).

PDR-1 colocalization with mitophagosomes was assessed in animals expressing the *foxSi30* (*pie-1p::DsRed::lgg-1::tbb-2 3’ UTR (oxti444)III*)*, fox60* (*3xFLAG::mNeonG::AID::pdr-1 III*) *and foxSi253* (*pie-1p::mts::TagBFP2::tbb-1 3’ UTR; (oxti179)II*) transgenes. Germlines of live animals were imaged using a Yokogawa spinning disk confocal microscope at 63× magnification, with images deconvolved as above. Colocalization events were scored using ImageJ. 3D renderings of events were generated using Imaris.

### DiOC6 (3,3′-dihexyloxacarbocyanine iodide) treatment

DiOC6 (3,3′-dihexyloxacarbocyanine iodide) was used as a mitochondrial membrane potential-sensitive dye^27^. NGM plates were supplemented with 500 nM of DiOC6 (Life Technologies), allowed to dry for 2 days, and then seeded with an OP50 *E. coli* culture containing 500 nM DiOC6 and allowed to sit for another 2 days. L4 animal were raised on the DiOC6 plates for 24 h at 20°C before imaging.

### Mitochondrial membrane potential quantification

Animals exposed to DiOC6 were imaged using a Yokogawa spinning disk confocal microscope at 63× magnification, as described above. Green (DiOC6), red (autophagosome), and blue (mitochondrial marker) channels were deconvolved prior to analysis. Image analysis was performed in Fiji/ImageJ using fixed intensity thresholds defined once and applied to all images.

### Lysotracker treatment

LysoTracker™ Deep Red (Invitrogen) was added to liquid NGM at a final concentration of 2 μM prior to plate pouring, and plates were allowed to solidify at room temperature. Plates were then seeded with OP50 *E. coli* bacteria supplemented with 2 μM LysoTracker™ Deep Red and incubated for 2 days to allow formation of a bacterial lawn containing the dye. Adult animals were killed by alkaline hypochlorite treatment and harvested eggs were transferred onto LysoTracker-containing NGM plates. Animals were maintained at 20°C and allowed to develop for ∼3–4 days until adulthood. Continuous exposure to LysoTracker™ Deep Red through both the agar and bacterial lawn enabled consistent labelling of acidic compartments prior to imaging.

### Autolysosome quantification

Autolysosomes in the germline were identified by colocalization of the autophagosome marker *foxSi30 (pie-1p::DsRed::lgg-1::tbb-2 3’ UTR (oxTi444)III)* with Lysotracker Deep Red, together with the mitochondrial marker *foxSi253 (pie-1p::mts::TagBFP2::tbb-2 3’ UTR (oxTi179)II)*. Germlines of live animals were imaged using a Yokogawa spinning disk confocal microscope at 63× magnification. Far-red (Lysotracker Deep Red), red (autophagosome), and blue (mitochondria) channels were deconvolved. Autolysosomes were defined as regions of overlap between the far-red and red signals. To quantify mitochondrial signal within these structures, a scan-line was manually drawn across individual autolysosomes, and fluorescence intensity profiles were obtained using the Plot Profile function in ImageJ. The mTagBFP2 signal along the profile was used to estimate fluorescence intensity within the autolysosomal region. Intensity and distance were normalized to their respective maxima (I/I^max^ and Dist/Dist^max^).

### Mitochondrial morphology quantification

L4-staged transgenic animals carrying *foxSi27* (*pie-1p::tomm-20::mKate2::HA::tbb-2 3’ UTR; (oxti385)I*) were imaged at 63× magnification using Yokogawa spinning disk confocal microscopy. A single z-plane was analyzed to quantify mitochondrial morphology by selecting the centremost z-plane. The image was first pre-processed in ImageJ to generate a binarized mask of germline mitochondria in the image. Briefly, the plugins ‘Subtract Background’, ‘Sigma Filter Plus’, ‘Enhance Local Contrast (CLAHE)’, ‘Adaptive Threshold’, and ‘Despeckle’ were used to pre-process the image, following a pipeline designed previously ^57^. The pre-processed image was then converted to a mask, and the area of each mitochondrial object was measured. The fraction of small mitochondria was calculated as the number of mitochondrial objects with an area < 0.2 µm^2^ divided by the total number of mitochondrial objects in the image. Secondly, the pre-processed image was skeletonized using ‘Skeletonize (2D/3D)’ in ImageJ to generate a branched network of the germline mitochondria. The skeletonized images were analysed for mean mitochondrial branch length using the plugin ‘MiNA Analyse Morphology’ ^58^. Alternatively, the plugin Mitochondria Analyzer (version 2.1.0), with default settings, was used to quantify mitochondrial morphology from different types of tissue. *fox43*(*hmg-5::Dendra2::FLAG*) *IV* and *foxsi253* (*pie-1p::mts::mTagBFP2::tbb-1 3’ UTR; (oxti179)II*) colocalization was quantified using ImageJ on images acquired by a Yokogawa spinning disk confocal microscopy and deconvolved as mentioned previously.

### mtDNA damage quantification

qPCR-based detection of mtDNA damage was performed following as previously described^39^. 20 animals were lysed in 50 μl of MLB, supplemented with 0.1 mg/ml proteinase K. The samples were lysed for 3 h at 65°C, and the proteinase subsequently heat-inactivated at 95°C for 15 min. qPCR was set up in 10 μl reactions with 5 μl of SensiFAST SYBR Green (Bioline), 0.5 μl of each primer and 1 μl of 1:100 diluted worm lysis. The PCR cycling conditions consisted of an initial denaturation step at 94°C for 5 min, followed by 40 cycles of 94°C for 10 s and 62°C for 45 s. qPCR was performed using a Rotor-Gene Q Real-Time PCR system. The primers nd-1f and nd-1r were used to amplify a short amplicon (75 bp) from mtDNA, and the primers ah113 and BLG3r were used to amplify the long amplicon (724 bp). Ct values were averaged across technical triplicates and analyzed using the ΔCt and ΔΔCt methods to calculate relative amplification (2^−ΔΔCt), with relative mtDNA damage estimated as −ln(2^-ΔΔCt^).

### Distal germ cell and unfertilized oocyte ΔmtDNA heteroplasmy

Single, 2-day-old animals carrying *spe-26(it112) IV* were transferred into 40 µl of M9 buffer supplemented with 2 nM of levamisole (Sigma) on a glass slide. Under a dissecting microscope, the animals were dissected using two 25 G needles. Entire gonads were separated from carcasses and unfertilized oocytes were further separated from the distal gonad containing germ cells. From the same individual, the distal germlines and 3 to 6 unfertilized oocytes were separated and then transferred into 3 µl of MLB^50^, supplemented with 0.1 mg/ml proteinase K. After lysis at 65°C for 1 h, qPCR was performed to quantify ΔmtDNA heteroplasmy as above.

A similar procedure was performed on 50 2-day-old animals using a microloader tip (Eppendorf). Here, 10 distal germlines were pooled and transferred to MLB, supplemented with 0.1 mg/ml proteinase K. The same procedure was performed for 10-20 unfertilized oocytes. After 1 h lysis at 65°C, qPCR was performed to quantify ΔmtDNA heteroplasmy. In these experiments, an independent data point represents the comparison between the average of three groups of 10 isolated distal germlines and the average of three groups of 10-20 isolated oocytes.

### Transgenerational inheritance of ΔmtDNA in mutant backgrounds

*uaDf5/+* hermaphrodites were crossed with mutant males. In F2 progeny, *uaDf5* heteroplasmy levels were measured by qPCR (as above) in either homozygous mutant or wild-type (control) nuclear backgrounds. F2 individuals with similar *uaDf5* heteroplasmy levels were selected and transferred to large (10 cm diameter) NGM plates seeded with OP50 *E. coli*. A large population of individuals was allowed to grow on the plate, and after 3 days, a 1 cm^2^ section of NGM was transferred to a new OP50-seeded plate for each generation, allowing the maintenance of the strain for ∼14 generations (F2 to F16) and the tracking of ΔmtDNA heteroplasmy. After cutting out the 1 cm^2^ slice, half of the plate was rinsed with M9 buffer and the worms were transferred to 100 of μl of MLB^50^, supplemented with 0.1 mg/ml proteinase K. The samples were lysed for 3 h at 65°C, and the proteinase subsequently heat-inactivated at 95°C for 15 min. Samples were diluted 1:200 in water and *uaDf5* levels determined as described above. At the end of experiment, for some conditions, 72 animals were collected and *uaDf5* levels determined individually as described above.

### Transgenerational inheritance of ΔmtDNA in transgenic backgrounds

A hermaphrodite animal deficient for mitophagy, carrying *pink-1(tm1779)*; *pdr-1(gk488)* or *dct-1(tm376)*, and a high and stable level of *uaDf5* heteroplasmy, was crossed with a transgenic male. F2 animals either transgenics or non-transgenics (control) were transferred to medium (5 cm diameter) NGM plates seeded with OP50. From F3, 30 larval stage 4 (L4) animals were transferred to a new medium NGM plate seeded with OP50, allowed to lay eggs for 2 days, before being collected for *uaDf5* heteroplasmy quantification by qPCR as described above. On the third day, 30 L4 animals were transferred to a new medium NGM plate seeded with OP50. This procedure was repeated for the following generations.

### Egg laying assay

L4 animals were separated onto individual small NGM plates seeded with OP50 and transferred to fresh plates every 24 h for 5–6 days; all plates were maintained at 20 °C. One day after removal of the adults, live progeny were counted on the plates.

### Mitochondrial affinity purification

Cell-specific mitochondrial affinity purification (CS-MAP) was performed as previously described^50,59^. Heteroplasmy in mitochondria was determined across large populations of animals (>10,000 individuals). This approach enabled detection of population-level trends by minimizing random inter-individual variability. Briefly, 10,000–20,000 worms consisting of a mixed population of late L4-stage and 1-day-old adults were grown on two 150 mm NGM plates seeded with *E. coli* OP50. Worms were homogenized using a hypotonic buffer^50^ and a Dounce homogenizer, and a crude mitochondrial fraction was subsequently enriched by differential centrifugation. HA-tagged baits were isolated from this fraction using anti-HA (influenza hemagglutinin) magnetic beads (Pierce). Washed beads were resuspended in 20 µl of hypotonic buffer. All CS-MAP experiments were repeated in at least three independent experiments. To control for population variability, purified mitochondria were compared to the corresponding total mitochondrial fraction (homogenate) from the same samples. Quantification of *uaDf5* heteroplasmy was performed as described previously.

### *C. elegans* Overall Fitness Index (COFI) quantification

A mixed population of animals from a large (10 cm diameter) NGM plate seeded with OP50 was collected in M9 and treated with sodium hypochlorite bleach to extract eggs. Eggs, resistant to the bleaching, were resuspended in 10 ml of M9 in a 15 ml conical tube and left to hatch for 24 h, under rotation, at room temperature. Newly hatched L1 were transferred to a large plate and single animals were separated onto individual small NGM plates seeded with OP50 on the next day. For 48 hours, animals were allowed to mature and lay eggs before they were removed from the plates. Forty-eight hours later, live progeny was scored. The live progeny scored reflects the ability of the parent animal to recover from the 24 h hatching and starvation step and reach the gravid stage, the ability of the parent animals to lay eggs, and the viability of the progeny.

### Human cell culture

The KSS cell line carrying the Kearns–Sayre syndrome (KSS)-associated mitochondrial DNA deletion (ΔmtDNA^4977^) was established by Moraes et al.^60^ and obtained from Professor Cole M. Haynes (University of Massachusetts, UMass)^61^. Stable transgenic cell lines were generated by lentiviral transduction of FLAG::BNIP3 under a TetOne doxycycline (Dox)-inducible promoter. Control and transgenic KSS cell lines were selected and expanded to reach heteroplasmy levels of ∼75–85% prior to experimentation. Cells were seeded in 6-well plates at 0.06 × 10⁶ cells per well and maintained at 37 °C with 5% CO₂ in DMEM high glucose (Gibco 11995-065), supplemented with 10% heat-inactivated FBS, 50 µg/ml uridine, and 1× antibiotic–antimycotic (Gibco 15240-062). Doxycycline was added at a final concentration of 0.5 μg/ml where indicated to induce transgene expression. Cells were passaged for multiple rounds under these conditions prior to reaching confluency. At each passage, a fraction of cells was harvested and lysed in buffer (50 mM Tris-HCl pH 7.4, 200 mM NaCl, 50 mM EDTA, 2% v/v SDS, 0.2 mg/ml proteinase K). Total DNA was extracted using phenol:chloroform:isoamyl alcohol (25:24:1, v/v/v).

### Quantification of ΔmtDNA^4977^ heteroplasmy

Quantitative PCR (qPCR) was performed using standard SYBR Green chemistry on a Rotor-Gene Q instrument with Rotor-Gene Q Pure Detection software (version 2.3.1, QIAGEN). Primer efficiencies were determined using a series of exponential dilutions (1 to 10⁻⁸ ng) of cloned wild-type human mtDNA (pSZ444) and ΔmtDNA^4977^ (pSZ430) fragments, enabling absolute copy number quantification. Primers AH582 and AH583 were used to quantify wild-type mtDNA levels, and AH580 and AH581 to quantify ΔmtDNA4977 levels (Supplementary Table 3). Cycling conditions were 95 °C for 2 min, followed by 40 cycles of 95 °C for 5 s, 58 °C for 10 s, and 72 °C for 15 s.

### Immunoblotting

Immunoblotting was performed as previously described^62^. Briefly, cells were harvested and lysed in SDS lysis buffer containing 50 mM Tris and 2% SDS by incubation at 97 °C for 15 min. Protein concentration was measured using a bicinchoninic acid (BCA) assay, and equal amounts of protein were separated by SDS–PAGE and transferred to membranes. FLAG::BNIP3 was detected using a rabbit anti-FLAG antibody (SAB4301135), and vinculin was detected using a mouse anti-vinculin antibody (sc-55465) as a loading control.

### Statistical analyses

For experiments comparing two samples, data were analyzed by an unpaired or paired two-way Student’s t-test. For experiments with more than two samples, data were analyzed using one-way ANOVA with Šídák’s multiple comparison test. Statistical tests were carried out using GraphPad Prism (version 11.0.0) and are annotated for each experiment in the corresponding figure legend. Biological replicates are represented on graphs

---

**Extended Data Fig. 1:**
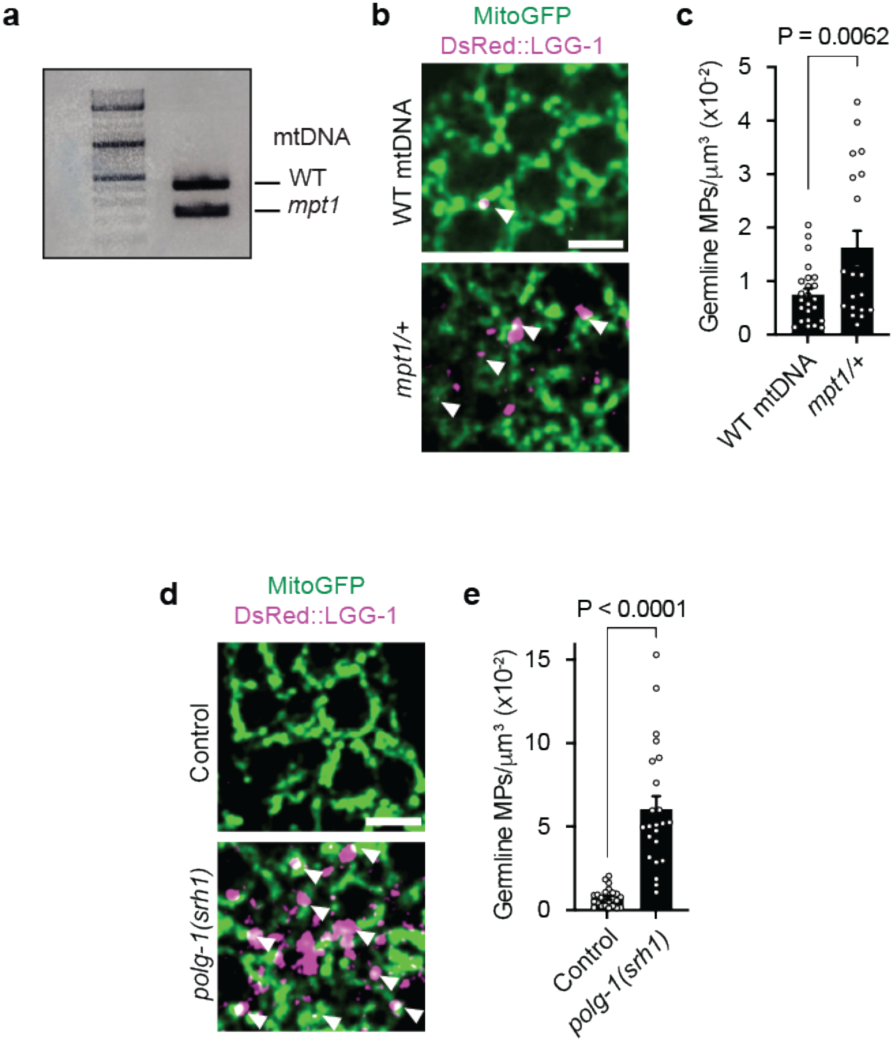
**a,** Semi-quantitative PCR analysis of *mpt1* heteroplasmy levels. DNA Ladder: O’GeneRuler DNA Ladder Mix. See Methods for details. **b,** Representative images of MPs in the germline in the presence or absence of *mpt1/+*. MPs are indicated with white arrowheads. Scale bar, 5 µm. **c,** Quantification of MPs normalized to mitochondrial volume in the germline in the presence or absence of *mpt1/+*. Columns are mean ± SEM; n = 24 (WT mtDNA) and 20 (*mpt1/+*) germlines per condition (3 independent experiments); unpaired two-tailed Student’s t-test. **d,** Representative images of MPs in the germline of control and *polg-1(srh1)* animals. MPs are indicated with white arrowheads. Scale bar, 5 µm. **e,** Quantification of MPs normalized to mitochondrial volume in the germline of control and *polg-1(srh1)* animals. Columns represent mean ± SEM; n = 24 (control) and 23 (*polg-1(srh1)*) germlines (3 independent experiments); unpaired two-tailed Student’s t-test.

**Extended Data Fig. 2:**
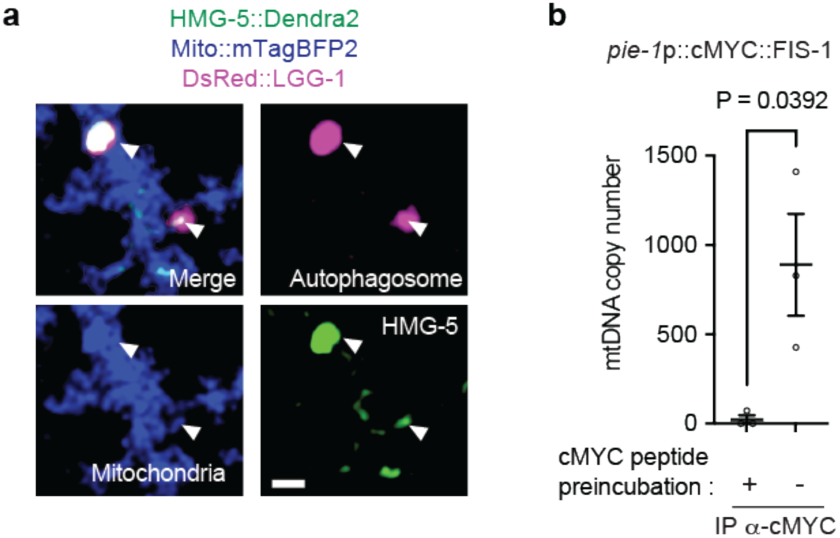
**a,** Representative image showing multiple MPs colocalizing with protein-mtDNA nucleoid complexes. Scale bar, 1 µm. **b,** qPCR analysis of immunoprecipitated mitochondria, purified from the germline, using germline-specific cMYC-tagged FIS-1 showing enrichment of mtDNA. For the control condition, anti-cMYC magnetic beads were pre-incubated with cMYC peptide. Bars represent mean ± SEM; n = 3 independent experiments; unpaired two-tailed Student’s t-test.

**Extended Data Fig. 3:**
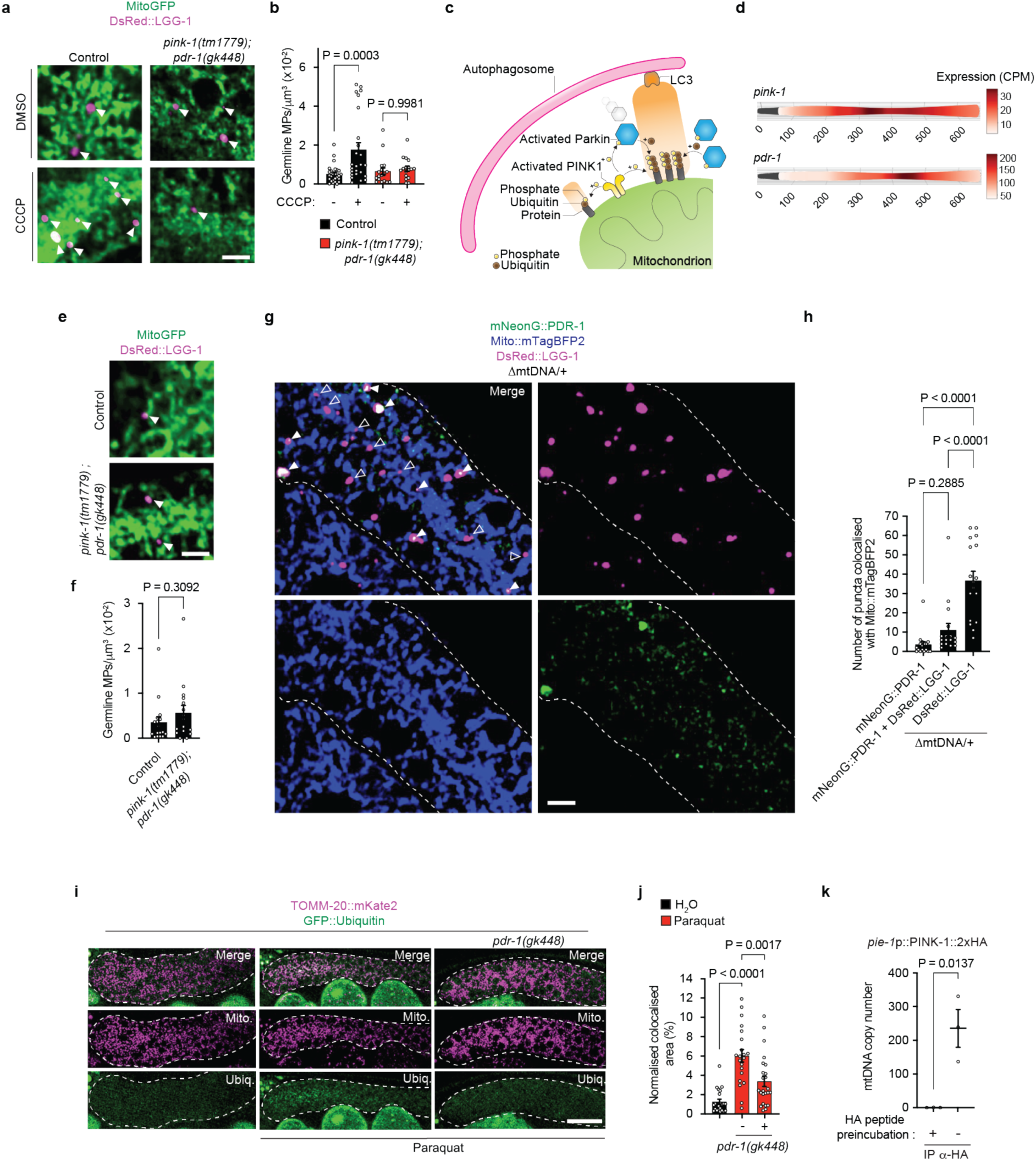
**a,** Representative images of MPs in the germline of control and *pink-1(tm1779);pdr-1(gk448)* mutant animals, treated with DMSO or CCCP. MPs are indicated with white arrowheads. Scale bar, 5 µm. **b,** Quantification of MPs normalized to mitochondrial volume in the germline of control and *pink-1(tm1779);pdr-1(gk448)* mutant animals, treated with DMSO or CCCP. Columns represent mean ± SEM; n = 27 (control, DMSO), 25 (control, CCCP), 17 (*pink-1(tm1779);pdr-1(gk448),* DMSO), and 17 (*pink-1(tm1779);pdr-1(gk448)*, CCCP) germlines (at least 3 independent experiments); one-way ANOVA with Šídák’s multiple comparisons test. **c,** Schematic of the PINK-1/Parkin mitophagy pathway. Stabilization of PINK-1 on the outer mitochondrial membrane leads to phosphorylation of pre-existing ubiquitin on mitochondrial proteins, initiating mitophagy signaling. PINK-1 also recruits and activates the E3 ubiquitin ligase Parkin through phosphorylation. Activated Parkin amplifies ubiquitination of mitochondrial substrates, reinforcing a feed-forward loop that promotes recruitment of LC3 and the formation of LC3-positive autophagosomes. **d,** Schematic of the spatial expression pattern of *pink-1* and *pdr-1* in the germline of *C. elegans*. Data are derived from the Spatial Caenorhabditis elegans Germline Expression of mRNA and miRNA (SPACEGERM)^63^. The y-axis represents expression levels (counts per million, CPM), and the x-axis indicates spatial coordinates corresponding to the physical distance (µm) along the distal-to-proximal axis of the gonad. **e,** Representative images of MPs in the germline of control and *pink-1(tm1779);pdr-1(gk448)* mutant animals. MPs are indicated with white arrowheads. Scale bar, 5 µm. **f,** Quantification of MPs normalized to mitochondrial volume in the germline of control and *pink-1(tm1779);pdr-1(gk448)* mutant animals. Columns represent mean ± SEM; n = 17 (control), 16 (*pink-1(tm1779);pdr-1(gk448)*). **g,** Representative images of MPs colocalizing with mNeonG::PDR-1 in the germline of animals carrying ΔmtDNA. MPs colocalizing with mNeonG::PDR-1 are indicated with solid white arrowheads, and non-colocalizing MPs with open white arrowheads. Scale bar, 5 µm. **h,** Quantification of colocalization with mitochondria for DsRed::LGG-1 alone, mNeonG::PDR-1 alone, and mNeonG::PDR-1 + DsRed::LGG-1 in the germline of animals carrying ΔmtDNA. Columns represent mean ± SEM; n = 17 germlines per condition (3 independent experiments); one-way ANOVA with Šídák’s multiple comparisons test. **i,** Representative images of GFP::Ubiquitin colocalization with TOMM-20::mKate2-labelled mitochondria in the germline of control and *pdr-1(gk448)* animals upon paraquat treatment. The germline is outlined with a white dashed line. Scale bar, 20 µm. **j,** Quantification of GFP::Ubiquitin and TOMM-20::mKate2 colocalization area in the germline of control and *pdr-1(gk448)* animals with or without paraquat treatment. Colocalization was quantified as total area using particle analysis ImageJ and normalized to germline area (arbitrary units). Columns are mean ± SEM; n = 22 (H_2_O), 21 (paraquat), and 24 (paraquat + *pdr-1(gk448)*) germlines (3 independent experiments); one-way ANOVA with Šídák’s multiple comparisons test. **k,** qPCR analysis of immunoprecipitated mitochondria, purified from the germline, using germline-specific PINK-1::2xHA showing enrichment of mtDNA. For the control condition, anti-HA magnetic beads were pre-incubated with HA peptide. Bars represent mean ± SEM; n = 3 independent experiments; unpaired two-tailed Student’s t-test.

**Extended Data Fig. 4:**
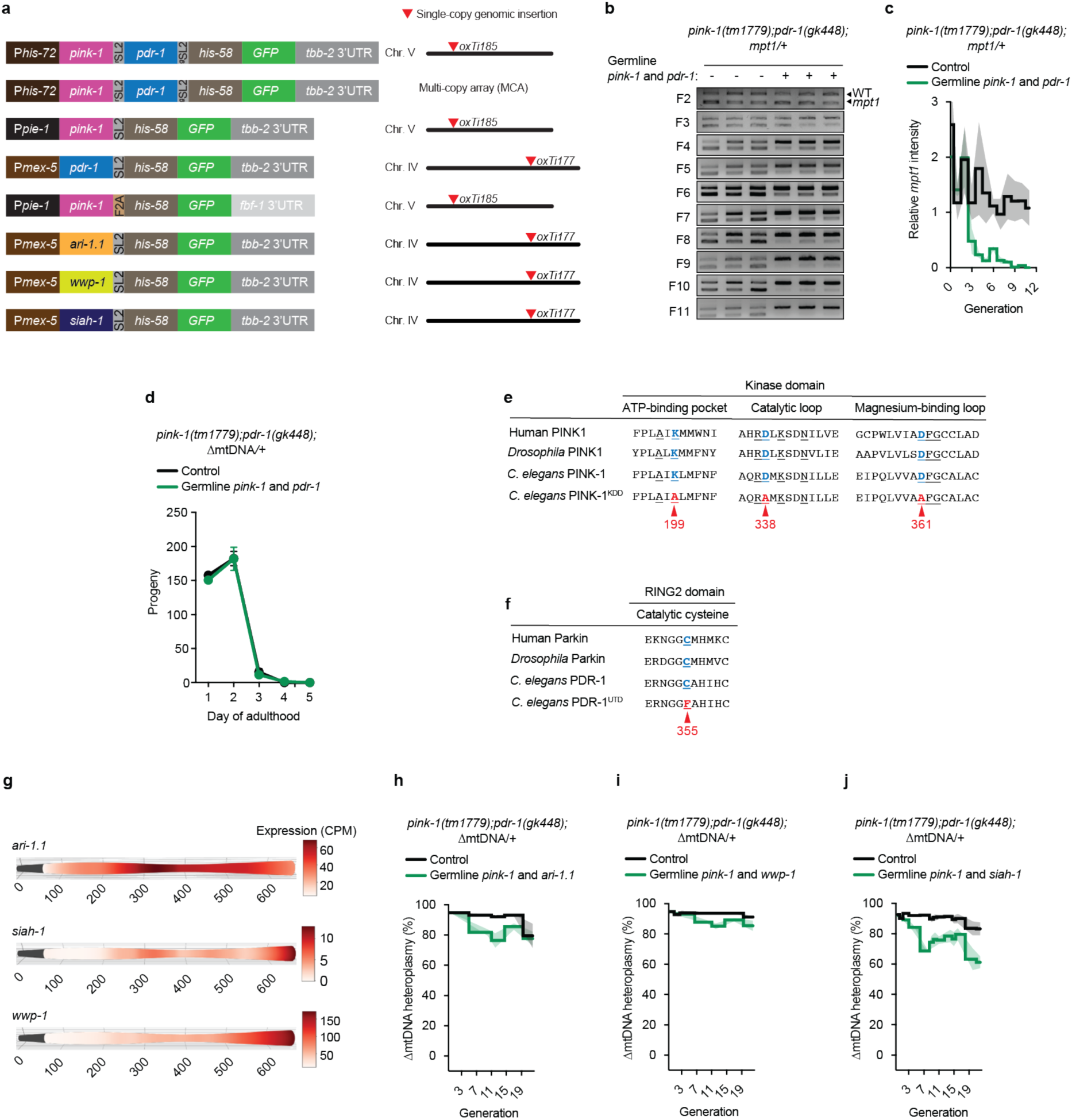
**a,** Schematics of some the transgenes used in this study and their genomic single-copy insertion sites. Single-copy insertion was performed using MosSCI^28^. The F2A peptide (Foot-and-mouth disease virus 2A) enables ribosomal skipping to generate two proteins from a single transcript^64^. The g/rSL2 sequence produces two independent mRNAs from a single transcript^65^. The *tbb-2 3′ UTR* is used for standard transgene expression, whereas the *fbf-1 3′ UTR* restricts expression to the distal germline aera of *C. elegans*^44^. **b,** Semi-quantitative PCR analysis of *mpt1* heteroplasmy across successive generations in *pink-1(tm1779);pdr-1(gk448);mpt1/+* animals, with or without germline expression of *pink-1* and *pdr-1*. 3 independent experiments are shown. **c,** Quantification of PCR band intensities from (**b**) using ImageJ. **d,** Egg-laying assay in *pink-1(tm1779);pdr-1(gk448);ΔmtDNA/+* animals with or without germline overexpression of *pink-1* and *pdr-1*. **e,** Generation of PINK-1(K199A,D338A,D361A), a kinase-dead (PINK-1^KDD^) triple mutant. ^66,67^. Human PINK1 (Q9BXM7), *Drosophila* PINK1 (Q0KHV6), and *C. elegans* PINK-1 (Q09298) were aligned using Clustal Omega (v1.2.4). Conserved residues within the kinase domain of *C. elegans* PINK-1 corresponding to the ATP-binding pocket (K199), catalytic loop (D338), and magnesium-binding loop (D361) were substituted with alanine (A). Underlined regions denote conserved amino acids; **blue** indicates wild-type residues, and **red** indicates substitutions, with position. **f,** Generation of a catalytically inactive PDR-1(C355F), a ubiquitin-transfer–defective (PDR-1^UTD^) mutant.^68^. Human Parkin (O60260), *Drosophila* Parkin (Q7KTX7), and *C. elegans* Parkin (Q9XUS3) were aligned using Clustal Omega (v1.2.4). Conserved catalytic cysteine (C355) within the RING2 domain (RBR E3 ligase) of *C. elegans* PDR-1 was substituted with phenylalanine **f,**. Conserved amino acid is underlined; **blue** indicates wild-type residue, and **red** indicates phenylalanine substitution, with position. **g,** Schematic diagram depicting the spatial expression pattern of *ari-1.1, siah-1,* and *wwp-1* in the germline of *C. elegans*. Data are derived from Spatial *Caenorhabditis elegans* germline expression of mRNA & miRNA (SPACEGERM)^63^. The y-axis represents expression levels (counts per million, CPM), and the x-axis indicates spatial coordinates corresponding to the physical distance (µm) along the distal-to-proximal axis of the gonad. (**h**–**j**) qPCR analysis of ΔmtDNA heteroplasmy across successive generations in *pink-1(tm1779);pdr-1(gk448);ΔmtDNA/+* animals overexpressing germline *pink-1* together with (**h**) germline *ari-1.1*, (**i**) germline *wwp-1*, or (**j**) germline *siah-1*. Data represent the mean of 3 independent experiments; shaded areas indicate ± SEM.

**Extended Data Fig. 5:**
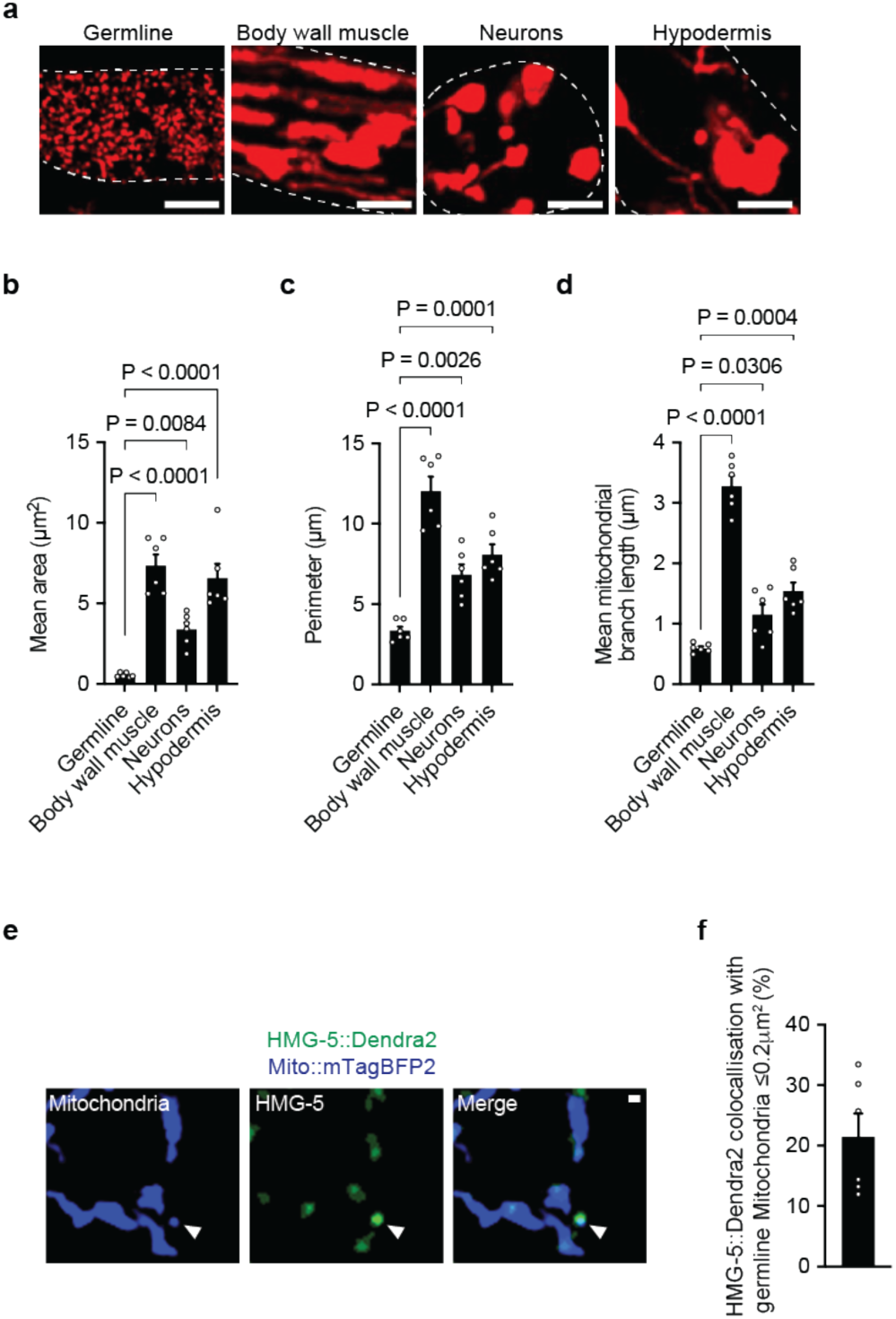
**a,** Representative images of mitochondria labelled with TOMM-20::mKate2::HA expressed specifically in the major tissues of *C. elegans*. Germline: *pie-1p::tomm-20::mKate2::HA::tbb-2 3’UTR],* body wall muscle: *myo-3p::tomm-20::mKate2::HA::tbb-2 3’UTR,* neurons: *rgef-1p::tomm-20::mKate2::HA::tbb-2 3’UTR*, and hypodermis: *dpy-7p::tomm-20::mKate2::HA::tbb-2 3’UTR*^59^. Tissue types are outlined with dashed white lines. Scale bar, 5 µm. (**b**–**d**) Quantification of (**b**) mean mitochondrial area, (**c**) perimeter, and (**d**) mean mitochondrial branch length in the germline, body wall muscle, neurons, and hypodermis. Columns are mean ± SEM; n = 6 independent animals per type of tissue; one-way ANOVA with Šídák’s multiple comparisons test. **e,** Representative images of colocalization of HMG-5::Dendra2 with mitochondria in the germline. Colocalization events are indicated with white arrowheads. Scale bar, 0.5 µm. **f,** Quantification of HMG-5::Dendra2 colocalization with ≤0.2 µm^2^ mitochondria in the germline. n = 6 independent animals.

**Extended Data Fig. 6:**
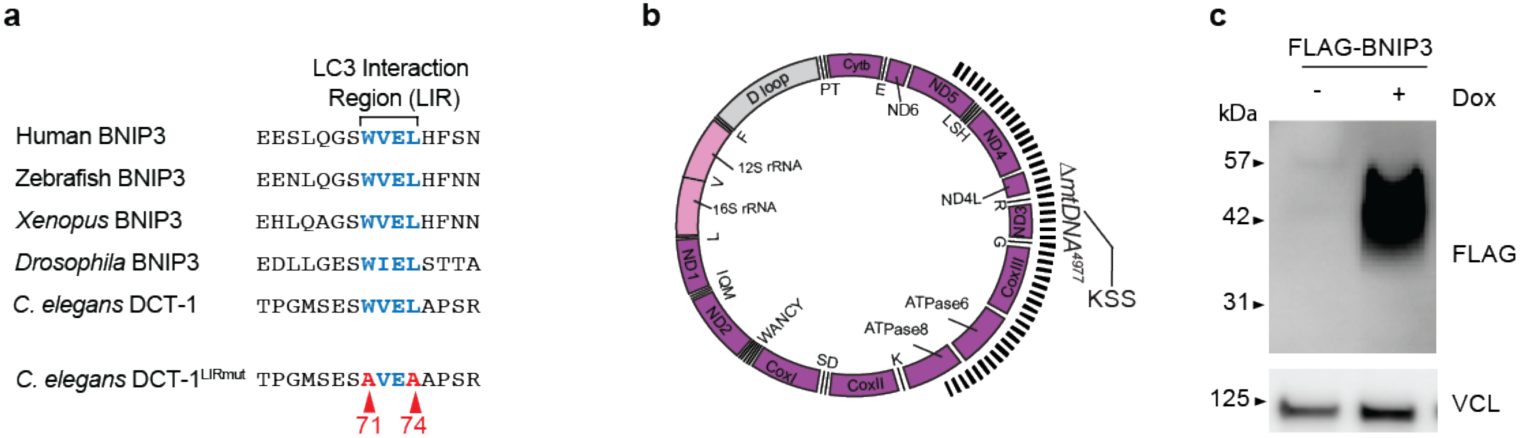
**a,** Generation of the DCT-1(W71A,L74A) (DCT-1^LIRmut^) mutant. BNIP3 sequences from human (Q12983), Zebrafish (Q5VK50), *Xenopus* (A0A8J0TCV6), *Drosophila* (Q9VPD6), and *C. elegans* (Q09969) were aligned using Clustal Omega (v1.2.4). The conserved LC3-interacting region (LIR) WVEL motif in *C. elegans* DCT-1 was mutated (W71A and L74A), generating an AVEA motif. **Blue** indicates wild-type residues, and **red** indicates alanine substitutions, with position. **b,** Schematic of human mtDNA showing the location of the Kearns–Sayre syndrome (KSS)-associated 4,977 bp deletion (*ΔmtDNA⁴⁹⁷⁷*). **c,** Representative western blot showing expression of FLAG–BNIP3 in KSS cells following doxycycline (Dox) induction. VCL, vinculin.

**Extended Data Fig. 7:**
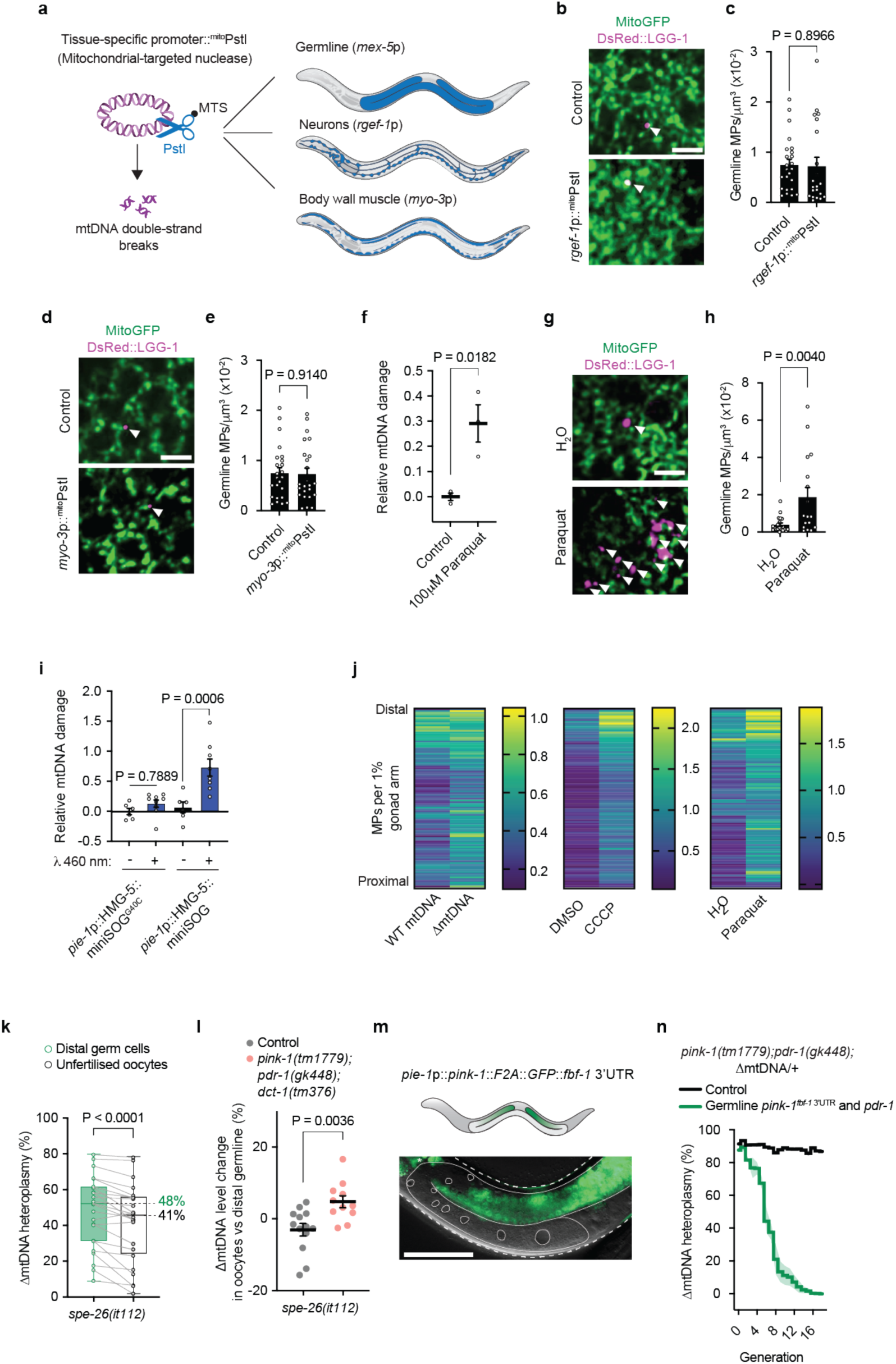
**a,** Schematic of the strategy used to induce tissue-specific mtDNA double-strand breaks (mtDSBs). ^MTS^PstI, a mitochondrially targeted endonuclease, induces mtDSBs by cleaving mtDNA. ^MTS^PstI expression was driven in a tissue-specific manner in the germline (*mex-5p*), neurons (*rgef-1p*), or body wall muscles (*myo-3p*). **b,** Representative images of MPs in the germline in the presence or absence of *rgef-1* p::^MTS^PstI. MPs are highlighted with white arrowheads. Scale bar, 5 µm. **c,** Quantification of MPs normalized to mitochondrial volume of the germline in the presence or absence of *rgef-1*p::^MTS^PstI. Columns represent mean ± SEM; n = 24 (control) and 21 (^MTS^PstI) germlines per condition (3 independent experiments); unpaired two-tailed Student’s t-test. **d,** Representative images of MPs in the germline in the presence or absence of *myo-3* p::^MTS^PstI. MPs are highlighted with white arrowheads. Scale bar, 5 µm. **e,** Quantification of MPs normalized to mitochondrial volume of the germline in the presence or absence of *myo-3*p::^MTS^PstI. Columns are mean ± SEM; n = 24 germlines per condition (3 independent experiments); unpaired two-tailed Student’s t-test. **f,** Quantification of relative mtDNA damage in control and animals treated with paraquat. Bars are mean ± SEM; n = 3 germlines per condition (3 independent experiments); unpaired two-tailed Student’s t-test. **g,** Representative images of MPs in the germline of animals treated with vehicle (H₂O) or paraquat. MPs are indicated with white arrowheads. Scale bar, 5 µm. **h,** Quantification of MPs normalized to mitochondrial volume in the germline of animals treated with vehicle (H₂O) or paraquat. Columns are mean ± SEM; n = 20 (H₂O) and 17 (paraquat) germlines (3 independent experiments); unpaired two-tailed Student’s t-test. **i,** Quantification of relative mtDNA damage in animals expressing *pie-1*p::HMG-5::miniSOG or the catalytically inactive variant *pie-1*p::HMG-5::miniSOG^G40C^, with or without 460 nm light photoactivation. Relative mtDNA damage was quantified by long/short amplicon PCR; reduced long-fragment amplification indicates increased lesion burden, including oxidative lesions and strand breaks^39^. Columns represent mean ± SEM; n = 6 (miniSOG^G40C^, non-illuminated), 9 (miniSOG^G40C^, illuminated), 6 (miniSOG, non-illuminated), and 8 (miniSOG, illuminated) germlines (3 independent experiments); one-way ANOVA with Šídák’s multiple comparisons test. **j,** Heat maps of individual mitophagy events along the gonad arm, normalized to gonad length from the distal to the proximal region. WT mtDNA vs ΔmtDNA/+; animals treated with DMSO or CCCP; animals treated with or without paraquat. Data are binned per 1% of gonad arm length; *n* = 100 animals. **k,** Quantification of ΔmtDNA levels in distal germ cells compared to unfertilized oocytes from the same dissected gonad in sperm-deficient *spe-26(it112)* animals. Data represent mean ± SEM; *n* = 24 independent gonads; paired two-tailed Student’s *t*-test. **l,** Quantification of ΔmtDNA levels in distal germ cells relative to unfertilized oocytes in control and *pink-1(tm1779);pdr-1(gk448);dct-1(tm376)* animals, both in a sperm-deficient *spe-26(it112)* background. Each data point represents the ratio of ΔmtDNA levels measured from ∼10 pooled distal germlines compared to ∼10–20 pooled unfertilized oocytes. Bars represent mean ± SEM; n = 13 (control) and 11 (triple mutant) independent experiments; unpaired two-tailed Student’s t-test. **m,** Representative image and schematic of the expression pattern of the *pie-1p::pink-1::F2A::his-58::GFP::fbf-1 3′UTR* transgene. The whole animal is outlined with a white dashed line, and the germline and oocyte nuclei are outlined with a solid white line. Scale bar, 50 µm. **n,** qPCR analysis of ΔmtDNA heteroplasmy across successive generations in *pink-1(tm1779); pdr-1(gk448)* animals carrying high levels of ΔmtDNA and in presence or absence of the *pie-1*p*::pink-1::F2A::his-58::GFP::fbf-1 3′UTR* (*pink-1*^fbf-1 3′ UTR^) together with *mex-5*p*::pdr-1*. Data represent the mean of 3 independent experiments; shaded areas indicate ± SEM.

## References

1. Jeedigunta, S.P., Minenkova, A.V., Palozzi, J.M., and Hurd, T.R. (2021). Avoiding Extinction: Recent Advances in Understanding Mechanisms of Mitochondrial DNA Purifying Selection in the Germline. Annu Rev Genomics Hum Genet 22, 55–80. 10.1146/annurev-genom-121420-081805.

2. Fan, W., Waymire, K.G., Narula, N., Li, P., Rocher, C., Coskun, P.E., Vannan, M.A., Narula, J., Macgregor, G.R., and Wallace, D.C. (2008). A mouse model of mitochondrial disease reveals germline selection against severe mtDNA mutations. Science 319, 958–962. 10.1126/science.1147786.

3. Floros, V.I., Pyle, A., Dietmann, S., Wei, W., Tang, W.W.C., Irie, N., Payne, B., Capalbo, A., Noli, L., Coxhead, J., et al. (2018). Segregation of mitochondrial DNA heteroplasmy through a developmental genetic bottleneck in human embryos. Nat Cell Biol 20, 144–151. 10.1038/s41556-017-0017-8.

4. Stewart, J.B., Freyer, C., Elson, J.L., and Larsson, N.G. (2008). Purifying selection of mtDNA and its implications for understanding evolution and mitochondrial disease. Nat Rev Genet 9, 657–662. 10.1038/nrg2396.

5. Wei, W., Tuna, S., Keogh, M.J., Smith, K.R., Aitman, T.J., Beales, P.L., Bennett, D.L., Gale, D.P., Bitner-Glindzicz, M.A.K., Black, G.C., et al. (2019). Germline selection shapes human mitochondrial DNA diversity. Science 364. 10.1126/science.aau6520.

6. Zaidi, A.A., Wilton, P.R., Su, M.S., Paul, I.M., Arbeithuber, B., Anthony, K., Nekrutenko, A., Nielsen, R., and Makova, K.D. (2019). Bottleneck and selection in the germline and maternal age influence transmission of mitochondrial DNA in human pedigrees. Proc Natl Acad Sci U S A 116, 25172–25178. 10.1073/pnas.1906331116.

7. Cree, L.M., Samuels, D.C., de Sousa Lopes, S.C., Rajasimha, H.K., Wonnapinij, P., Mann, J.R., Dahl, H.H., and Chinnery, P.F. (2008). A reduction of mitochondrial DNA molecules during embryogenesis explains the rapid segregation of genotypes. Nat Genet 40, 249–254. 10.1038/ng.2007.63.

8. Jenuth, J.P., Peterson, A.C., Fu, K., and Shoubridge, E.A. (1996). Random genetic drift in the female germline explains the rapid segregation of mammalian mitochondrial DNA. Nat Genet 14, 146–151. 10.1038/ng1096-146.

9. Wai, T., Teoli, D., and Shoubridge, E.A. (2008). The mitochondrial DNA genetic bottleneck results from replication of a subpopulation of genomes. Nat Genet 40, 1484–1488. 10.1038/ng.258.

10. Zhang, Y., Wang, Z.H., Liu, Y., Chen, Y., Sun, N., Gucek, M., Zhang, F., and Xu, H. (2019). PINK1 Inhibits Local Protein Synthesis to Limit Transmission of Deleterious Mitochondrial DNA Mutations. Mol Cell 73, 1127–1137 e1125. 10.1016/j.molcel.2019.01.013.

11. Hill, J.H., Chen, Z., and Xu, H. (2014). Selective propagation of functional mitochondrial DNA during oogenesis restricts the transmission of a deleterious mitochondrial variant. Nat Genet. ng.2920 [pii] 10.1038/ng.2920.

12. Lieber, T., Jeedigunta, S.P., Palozzi, J.M., Lehmann, R., and Hurd, T.R. (2019). Mitochondrial fragmentation drives selective removal of deleterious mtDNA in the germline. Nature. 10.1038/s41586-019-1213-4.

13. Ma, H., Xu, H., and O’Farrell, P.H. (2014). Transmission of mitochondrial mutations and action of purifying selection in Drosophila melanogaster. Nat Genet. ng.2919 [pii] 10.1038/ng.2919.

14. Gao, Q., Tian, R., Han, H., Slone, J., Wang, C., Ke, X., Zhang, T., Li, X., He, Y., Liao, P., et al. (2022). PINK1-mediated Drp1(S616) phosphorylation modulates synaptic development and plasticity via promoting mitochondrial fission. Signal Transduct Target Ther 7, 103. 10.1038/s41392-022-00933-z.

15. Huang, E., Qu, D., Huang, T., Rizzi, N., Boonying, W., Krolak, D., Ciana, P., Woulfe, J., Klein, C., Slack, R.S., et al. (2017). PINK1-mediated phosphorylation of LETM1 regulates mitochondrial calcium transport and protects neurons against mitochondrial stress. Nat Commun 8, 1399. 10.1038/s41467-017-01435-1.

16. Matheoud, D., Sugiura, A., Bellemare-Pelletier, A., Laplante, A., Rondeau, C., Chemali, M., Fazel, A., Bergeron, J.J., Trudeau, L.E., Burelle, Y., et al. (2016). Parkinson’s Disease-Related Proteins PINK1 and Parkin Repress Mitochondrial Antigen Presentation. Cell 166, 314–327. 10.1016/j.cell.2016.05.039.

17. McLelland, G.L., Soubannier, V., Chen, C.X., McBride, H.M., and Fon, E.A. (2014). Parkin and PINK1 function in a vesicular trafficking pathway regulating mitochondrial quality control. EMBO J 33, 282–295. 10.1002/embj.201385902.

18. Narendra, D.P., Jin, S.M., Tanaka, A., Suen, D.F., Gautier, C.A., Shen, J., Cookson, M.R., and Youle, R.J. (2010). PINK1 Is Selectively Stabilized on Impaired Mitochondria to Activate Parkin. Plos Biology 8. ARTN e1000298 10.1371/journal.pbio.1000298.

19. Schwartz, A.Z.A., Tsyba, N., Abdu, Y., Patel, M.R., and Nance, J. (2022). Independent regulation of mitochondrial DNA quantity and quality in Caenorhabditis elegans primordial germ cells. Elife 11. 10.7554/eLife.80396.

20. Kremer, L.S., Bozhilova, L.V., Rubalcava-Gracia, D., Filograna, R., Upadhyay, M., Koolmeister, C., Chinnery, P.F., and Larsson, N.G. (2023). A role for BCL2L13 and autophagy in germline purifying selection of mtDNA. PLoS Genet 19, e1010573. 10.1371/journal.pgen.1010573.

21. Palozzi, J.M., Jeedigunta, S.P., Minenkova, A.V., Monteiro, V.L., Thompson, Z.S., Lieber, T., and Hurd, T.R. (2022). Mitochondrial DNA quality control in the female germline requires a unique programmed mitophagy. Cell Metab 34, 1809–1823 e1806. 10.1016/j.cmet.2022.10.005.

22. Thendral, S.B., Bacot, S., Ryde, I.T., Morton, K.S., Chi, Q., Kenny-Ganzert, I.W., Meyer, J.N., and Sherwood, D.R. (2026). Programmed mitophagy at the oocyte-to-zygote transition promotes lineage endurance. Nat Cell Biol. 10.1038/s41556-025-01854-z.

23. Frison, M., Lockey, B.S., Nie, Y., Golder, Z., Theiaspra, E., Ryall, C.D., Lyons, C., Burr, S.P., Prater, M., Bozhilova, L.V., et al. (2025). Ubiquitin-mediated mitophagy regulates the inheritance of mitochondrial DNA mutations. Science 390, 156–163. 10.1126/science.adr5438.

24. Palikaras, K., Lionaki, E., and Tavernarakis, N. (2015). Coordination of mitophagy and mitochondrial biogenesis during ageing in C. elegans. Nature 521, 525–528. 10.1038/nature14300.

25. Haroon, S., Li, A., Weinert, J.L., Fritsch, C., Ericson, N.G., Alexander-Floyd, J., Braeckman, B.P., Haynes, C.M., Bielas, J.H., Gidalevitz, T., and Vermulst, M. (2018). Multiple molecular mechanisms rescue mtDNA disease in *C. elegans*. Cell Rep 22, 3115–3125. 10.1016/j.celrep.2018.02.099.

26. Ahier, A., Dai, C.-Y., Tweedie, A., Bezawork-Geleta, A., Kirmes, I., and Zuryn, S. (2018). Affinity purification of cell-specific mitochondria from whole animals resolves patterns of genetic mosaicism. Nature Cell Biology 20, 352–360. doi: 10.1038/s41556-017-0023-x.

27. Bohnert, K.A., and Kenyon, C. (2017). A lysosomal switch triggers proteostasis renewal in the immortal *C. elegans* germ lineage. Nature 551, 629–633. 10.1038/nature24620.

28. Frokjaer-Jensen, C., Davis, M.W., Sarov, M., Taylor, J., Flibotte, S., Labella, M., Pozniakovsky, A., Moerman, D.G., and Jorgensen, E.M. (2014). Random and targeted transgene insertion in Caenorhabditis elegans using a modified Mos1 transposon. Nat Methods 11, 529–534. nmeth.2889 [pii] 10.1038/nmeth.2889.

29. Kelly, W.G., Xu, S., Montgomery, M.K., and Fire, A. (1997). Distinct requirements for somatic and germline expression of a generally expressed Caernorhabditis elegans gene. Genetics 146, 227–238. 10.1093/genetics/146.1.227.

30. Kleele, T., Rey, T., Winter, J., Zaganelli, S., Mahecic, D., Perreten Lambert, H., Ruberto, F.P., Nemir, M., Wai, T., Pedrazzini, T., and Manley, S. (2021). Distinct fission signatures predict mitochondrial degradation or biogenesis. Nature 593, 435–439. 10.1038/s41586-021-03510-6.

31. Breckenridge, D.G., Kang, B.H., Kokel, D., Mitani, S., Staehelin, L.A., and Xue, D. (2008). Caenorhabditis elegans drp-1 and fis-2 regulate distinct cell-death execution pathways downstream of ced-3 and independent of ced-9. Mol Cell 31, 586–597. 10.1016/j.molcel.2008.07.015.

32. Campbell, D., and Zuryn, S. (2023). The mechanisms and roles of mitochondrial dynamics in C. elegans. Semin Cell Dev Biol. 10.1016/j.semcdb.2023.10.006.

33. Zhu, Y., Massen, S., Terenzio, M., Lang, V., Chen-Lindner, S., Eils, R., Novak, I., Dikic, I., Hamacher-Brady, A., and Brady, N.R. (2013). Modulation of serines 17 and 24 in the LC3-interacting region of Bnip3 determines pro-survival mitophagy versus apoptosis. J Biol Chem 288, 1099–1113. 10.1074/jbc.M112.399345.

34. Pickles, S., Vigie, P., and Youle, R.J. (2018). Mitophagy and Quality Control Mechanisms in Mitochondrial Maintenance. Curr Biol 28, R170–R185. 10.1016/j.cub.2018.01.004.

35. He, Y.L., Li, J., Gong, S.H., Cheng, X., Zhao, M., Cao, Y., Zhao, T., Zhao, Y.Q., Fan, M., Wu, H.T., et al. (2022). BNIP3 phosphorylation by JNK1/2 promotes mitophagy via enhancing its stability under hypoxia. Cell Death Dis 13, 966. 10.1038/s41419-022-05418-z.

36. Palikaras, K., Lionaki, E., and Tavernarakis, N. (2015). Coupling mitogenesis and mitophagy for longevity. Autophagy 11, 1428–1430. 10.1080/15548627.2015.1061172.

37. Zhang, T., Xue, L., Li, L., Tang, C., Wan, Z., Wang, R., Tan, J., Tan, Y., Han, H., Tian, R., et al. (2016). BNIP3 Protein Suppresses PINK1 Kinase Proteolytic Cleavage to Promote Mitophagy. J Biol Chem 291, 21616–21629. 10.1074/jbc.M116.733410.

38. Wang, S., Long, H., Hou, L., Feng, B., Ma, Z., Wu, Y., Zeng, Y., Cai, J., Zhang, D.W., and Zhao, G. (2023). The mitophagy pathway and its implications in human diseases. Signal Transduct Target Ther 8, 304. 10.1038/s41392-023-01503-7.

39. Dai, C.Y., Ng, C.C., Hung, G.C.C., Kirmes, I., Hughes, L.A., Du, Y., Brosnan, C.A., Ahier, A., Hahn, A., Haynes, C.M., et al. (2023). ATFS-1 counteracts mitochondrial DNA damage by promoting repair over transcription. Nature Cell Biology 25, 1111–1120. 10.1038/s41556-023-01192-y.

40. Li, Y., and Cui, Z.J. (2020). Photodynamic Activation of Cholecystokinin 1 Receptor with Different Genetically Encoded Protein Photosensitizers and from Varied Subcellular Sites. Biomolecules 10. 10.3390/biom10101423.

41. Kukat, C., Wurm, C.A., Spahr, H., Falkenberg, M., Larsson, N.G., and Jakobs, S. (2011). Super-resolution microscopy reveals that mammalian mitochondrial nucleoids have a uniform size and frequently contain a single copy of mtDNA. Proc Natl Acad Sci U S A 108, 13534–13539. 1109263108 [pii] 10.1073/pnas.1109263108.

42. Varkey, J.P., Muhlrad, P.J., Minniti, A.N., Do, B., and Ward, S. (1995). The Caenorhabditis elegans spe-26 gene is necessary to form spermatids and encodes a protein similar to the actin-associated proteins kelch and scruin. Genes Dev 9, 1074–1086. 10.1101/gad.9.9.1074.

43. Merritt, C., Rasoloson, D., Ko, D., and Seydoux, G. (2008). 3’ UTRs are the primary regulators of gene expression in the C. elegans germline. Curr Biol 18, 1476–1482. 10.1016/j.cub.2008.08.013.

44. Merritt, C., and Seydoux, G. (2010). Transgenic solutions for the germline. WormBook, 1–21. 10.1895/wormbook.1.148.1.

45. Charmpilas, N., and Tavernarakis, N. (2020). Mitochondrial maturation drives germline stem cell differentiation in Caenorhabditis elegans. Cell Death Differ 27, 601–617. 10.1038/s41418-019-0375-9.

46. Suen, D.F., Narendra, D.P., Tanaka, A., Manfredi, G., and Youle, R.J. (2010). Parkin overexpression selects against a deleterious mtDNA mutation in heteroplasmic cybrid cells. Proc Natl Acad Sci U S A 107, 11835–11840. 10.1073/pnas.0914569107.

47. Brenner, S. (1974). The genetics of Caenorhabditis elegans. Genetics 77, 71–94.

48. Silva-Garcia, C.G., Lanjuin, A., Heintz, C., Dutta, S., Clark, N.M., and Mair, W.B. (2019). Single-Copy Knock-In Loci for Defined Gene Expression in Caenorhabditis elegans. G3 (Bethesda) 9, 2195–2198. 10.1534/g3.119.400314.

49. Gibson, D.G., Young, L., Chuang, R.-Y., Venter, J.C., Hutchison, C.A., and Smith, H.O. (2009). Enzymatic assembly of DNA molecules up to several hundred kilobases. Nat Methods 6, 343–345.

50. Ahier, A., Onraet, T., and Zuryn, S. (2021). Cell-specific mitochondria affinity purification (CS-MAP) from Caenorhabditis elegans. STAR Protoc 2, 100952. 10.1016/j.xpro.2021.100952.

51. Miyazaki, K. (2011). MEGAWHOP cloning: a method of creating random mutagenesis libraries via megaprimer PCR of whole plasmids. Methods Enzymol 498, 399–406. 10.1016/B978-0-12-385120-8.00017-6.

52. Noma, K., and Jin, Y. (2015). Optogenetic mutagenesis in Caenorhabditis elegans. Nat Commun 6, 8868. 10.1038/ncomms9868.

53. Ashley, G.E., Duong, T., Levenson, M.T., Martinez, M.A.Q., Johnson, L.C., Hibshman, J.D., Saeger, H.N., Palmisano, N.J., Doonan, R., Martinez-Mendez, R., et al. (2021). An expanded auxin-inducible degron toolkit for Caenorhabditis elegans. Genetics 217. 10.1093/genetics/iyab006.

54. Gitschlag, B.L., Kirby, C.S., Samuels, D.C., Gangula, R.D., Mallal, S.A., and Patel, M.R. (2016). Homeostatic responses regulate selfish mitochondrial genome dynamics in *C. elegans*. Cell Metab 24, 91–103. 10.1016/j.cmet.2016.06.008.

55. Schindelin, J., Arganda-Carreras, I., Frise, E., Kaynig, V., Longair, M., Pietzsch, T., Preibisch, S., Rueden, C., Saalfeld, S., Schmid, B., et al. (2012). Fiji: an open-source platform for biological-image analysis. Nat Methods 9, 676–682. 10.1038/nmeth.2019.

56. Palikaras, K., and Tavernarakis, N. (2017). In vivo Mitophagy Monitoring in Caenorhabditis elegans to Determine Mitochondrial Homeostasis. Bio Protoc 7. 10.21769/BioProtoc.2215.

57. Chaudhry, A., Shi, R., and Luciani, D.S. (2020). A pipeline for multidimensional confocal analysis of mitochondrial morphology, function, and dynamics in pancreatic β-cells. Am J Physiol Endocrinol Metab 318, E87–e101. 10.1152/ajpendo.00457.2019.

58. Valente, A.J., Maddalena, L.A., Robb, E.L., Moradi, F., and Stuart, J.A. (2017). A simple ImageJ macro tool for analyzing mitochondrial network morphology in mammalian cell culture. Acta Histochem 119, 315–326. 10.1016/j.acthis.2017.03.001.

59. Ahier, A., Dai, C.Y., Tweedie, A., Bezawork-Geleta, A., Kirmes, I., and Zuryn, S. (2018). Publisher Correction: Affinity purification of cell-specific mitochondria from whole animals resolves patterns of genetic mosaicism. Nat Cell Biol 20, 361. 10.1038/s41556-018-0055-x.

60. Moraes, C.T., DiMauro, S., Zeviani, M., Lombes, A., Shanske, S., Miranda, A.F., Nakase, H., Bonilla, E., Werneck, L.C., Servidei, S., and, et al. (1989). Mitochondrial DNA deletions in progressive external ophthalmoplegia and Kearns-Sayre syndrome. N Engl J Med 320, 1293–1299. 10.1056/NEJM198905183202001.

61. Yang, Q.Y., Liu, P.P., Anderson, N.S., Shpilka, T., Du, Y.G., Naresh, N.U., Li, R., Zhu, L.J., Luk, K., Lavelle, J., et al. (2022). LONP-1 and ATFS-1 sustain deleterious heteroplasmy by promoting mtDNA replication in dysfunctional mitochondria. Nature Cell Biology 24, 181-+. 10.1038/s41556-021-00840-5.

62. Nguyen-Dien, G.T., Kozul, K.L., Cui, Y., Townsend, B., Kulkarni, P.G., Ooi, S.S., Marzio, A., Carrodus, N., Zuryn, S., Pagano, M., et al. (2023). FBXL4 suppresses mitophagy by restricting the accumulation of NIX and BNIP3 mitophagy receptors. EMBO J 42, e112767. 10.15252/embj.2022112767.

63. Diag, A., Schilling, M., Klironomos, F., Ayoub, S., and Rajewsky, N. (2018). Spatiotemporal m(i)RNA Architecture and 3’ UTR Regulation in the C. elegans Germline. Dev Cell 47, 785–800 e788. 10.1016/j.devcel.2018.10.005.

64. Ahier, A., and Jarriault, S. (2014). Simultaneous expression of multiple proteins under a single promoter in Caenorhabditis elegans via a versatile 2A-based toolkit. Genetics 196, 605–613. 10.1534/genetics.113.160846.

65. Spieth, J., Brooke, G., Kuersten, S., Lea, K., and Blumenthal, T. (1993). Operons in C. elegans: polycistronic mRNA precursors are processed by trans-splicing of SL2 to downstream coding regions. Cell 73, 521–532. 10.1016/0092-8674(93)90139-h.

66. Beilina, A., Van Der Brug, M., Ahmad, R., Kesavapany, S., Miller, D.W., Petsko, G.A., and Cookson, M.R. (2005). Mutations in PTEN-induced putative kinase 1 associated with recessive parkinsonism have differential effects on protein stability. Proc Natl Acad Sci U S A 102, 5703–5708. 10.1073/pnas.0500617102.

67. Pridgeon, J.W., Olzmann, J.A., Chin, L.S., and Li, L. (2007). PINK1 protects against oxidative stress by phosphorylating mitochondrial chaperone TRAP1. PLoS Biol 5, e172. 10.1371/journal.pbio.0050172.

68. Lazarou, M., Narendra, D.P., Jin, S.M., Tekle, E., Banerjee, S., and Youle, R.J. (2013). PINK1 drives Parkin self-association and HECT-like E3 activity upstream of mitochondrial binding. J Cell Biol 200, 163–172. 10.1083/jcb.201210111.

